# The global distribution of angiosperm genome size is shaped by climate

**DOI:** 10.1101/2022.12.05.519116

**Authors:** Petr Bureš, Tammy L. Elliott, Pavel Veselý, Petr Šmarda, Félix Forest, Ilia J. Leitch, Eimear Nic Lughadha, Marybel Soto Gomez, Samuel Pironon, Matilda J. M. Brown, Jakub Šmerda, František Zedek

## Abstract

1. Angiosperms, which inhabit diverse environments across all continents, exhibit significant variation in genome sizes, making them an excellent model system for examining hypotheses about the global distribution of genome size. These include the previously proposed large-genome-constraint, mutational-hazard, polyploidy-mediated, and climate-mediated hypotheses.
2. We compiled the largest genome size dataset to date, encompassing >5% of known angiosperm species, and analyzed genome size distribution using a comprehensive geographic distribution dataset for all angiosperms.
3. We observed that angiosperms with large range sizes generally had small genomes, supporting the large-genome-constraint hypothesis. Climate was shown to exert a strong influence on genome size distribution along the global latitudinal gradient, while the frequency of polyploidy and the type of growth form had negligible effects. In contrast to the unimodal patterns along the global latitudinal gradient shown by plant size traits and polyploid proportions, the increase in angiosperm genome size from the equator to 40-50°N/S is probably mediated by different (mostly climatic) mechanisms than the decrease in genome sizes observed from 40–50°N northwards.
4. Our analysis suggests that the global distribution of genome sizes in angiosperms is mainly shaped by climatically-mediated purifying selection, genetic drift, relaxed selection, and environmental filtering.

## Introduction

The most essential structure of any organism is its genome, of which the size is a relatively stable species-specific property. Angiosperms exhibit tremendous variation in genome sizes (more than 2,400-fold; Pellicer *et al*., 2018) and are found across all continents, with the majority of species being narrow endemics while a minority are widespread cosmopolitan species (Enquist *et al*., 2019). This makes angiosperms a powerful model system for studying the underlying drivers that shape genome size evolution and its distribution across the globe. The recent increase in the use of flow cytometry in botanical studies has led to a substantial accumulation of standardized genome size data across wide phylogenetic and geographic scales (Garcia *et al*., 2014; Leitch *et al*., 2019; Šmarda *et al*., 2019; Zonneveld, 2019). Given that consistent geographic data has recently become available for most known species through the World Checklist of Vascular Plants (WCVP; Govaerts *et al*., 2021), it is now possible to examine hypotheses seeking to understand the causal links between angiosperm genome size, distribution, and environment at a global scale.

Key proximal mechanisms generating changes in genome size are polyploidization followed by re-diploidization (Wendel, 2000; Leitch & Leitch, 2008; Soltis *et al*., 2015; Guignard *et al*., 2016; Šmarda *et al*., 2019) and the accumulation and removal of repetitive DNA (Levin, 2002; Wendel *et al*., 2016; Lwin *et al*., 2017), especially transposable elements (TEs), which constitute the main component of most plant genomes (Bennetzen *et al*., 2005; Tenaillon *et al*., 2010; Lisch, 2013; Bennetzen & Wang, 2014).

The ’large-genome-constraint’ hypothesis (LGCH) suggests that species with large genomes might face selection pressure against them due to their negative impact on plant anatomy and physiology (Vinogradov, 2003; Knight *et al*., 2005). This is because more genomic material occupies a larger volume, influencing the minimum cell size (Cavalier-Smith, 2005; Šímová & Herben, 2012; Bhadra *et al*., 2023). Consequently, plants with larger genomes tend to have larger seeds (Knight & Ackerly, 2002; Beaulieu *et al*., 2007; Carta *et al*., 2022; Bhadra *et al*., 2023), a trait linked to smaller distributional ranges (Sonkoly *et al*., 2022). Additionally, they possess larger stomatal guard cells (Beaulieu et al., 2008; Veselý *et al*., 2012; Bhadra *et al*., 2023), which close and open more slowly (Drake et al., 2013; Kardiman & Ræbild, 2018; Lawson & Matthews, 2020). This might be disadvantageous in, for example, arid environments that demand efficient water management (Veselý *et al*., 2020; Bureš *et al*., 2023; Šmarda *et al*., 2023). Larger cells also limit the mesophyll surface area packed into the leaf volume leading to lower CO_2_ diffusion and rates of photosynthesis (Théroux-Rancourt *et al*., 2021). Species with large genomes also experience slower rates of cell division (Francis *et al*., 2008; Šímová & Herben, 2012) and have higher phosphorus (P) and/or nitrogen (N) requirements (Šmarda *et al*., 2013; Peng *et al*., 2022). Large genomes may thus limit species’ dispersal abilities and have narrower ecological niches, potentially resulting in smaller geographic ranges (Sheth *et al*., 2020). In contrast, smaller genomes offer more flexibility in cell size (Beaulieu *et al*., 2007; Beaulieu *et al*., 2008; Veselý *et al*., 2012; Meyerson *et al*., 2020; Bhadra *et al*., 2023), have faster rates of cell division (Francis *et al*., 2008; Šímová & Herben, 2012), and lower P and N demands (Šmarda *et al*., 2013; Peng *et al*., 2022) allowing greater plasticity in range size.

Although TE insertions can occasionally have adaptive effects (Casacuberta & González, 2013; Schrader & Schmitz, 2019), they are mostly neutral or deleterious (Deniz *et al*., 2019). Thus, TE insertions mostly become fixed via genetic drift rather than by natural selection or intragenomic selection favoring TE accumulation (Werren, 2011; Deniz *et al*., 2019). As the relative importance of natural selection versus random genetic drift depends on population size, the mutational-hazard hypothesis (MHH) posits that genome growth via TEs occurs more readily in smaller populations, where genetic drift is more prominent than natural selection (i.e., species with smaller effective population sizes will have larger genomes; Lynch & Conery, 2003; Lynch, 2007). The relative importance of natural selection and genetic drift also appears to hold for species range size in both plants and animals (Corbett-Detig *et al*., 2015), likely because of the positive abundance-occupation relationship (Gaston *et al*., 2002) where species with larger populations tend to have large distributional ranges (e.g., Brown, 1984; Johnson, 1998; Gaston, 2003; Webb *et al*., 2012; Drovetski *et al*., 2014; Spence *et al*., 2021; Guo *et al*., 2022; Ten Caten *et al*., 2022).

Considering the potential effects of genetic drift and natural selection on genome size and their interplay with range size, the LGCH predicts that species with large ranges should not have large genomes, resulting in a triangular relationship (Fig. 1a). On the other hand, the MHH predicts that species genome sizes should decrease with increasing geographic ranges, producing a negative relationship (Fig. 1b). Although effective population size is affected by complex factors and range size is a relatively crude proxy, the high statistical power provided by the large amount of currently available data on species genome size and distribution should help overcome this imprecision.

**Fig. 1.**
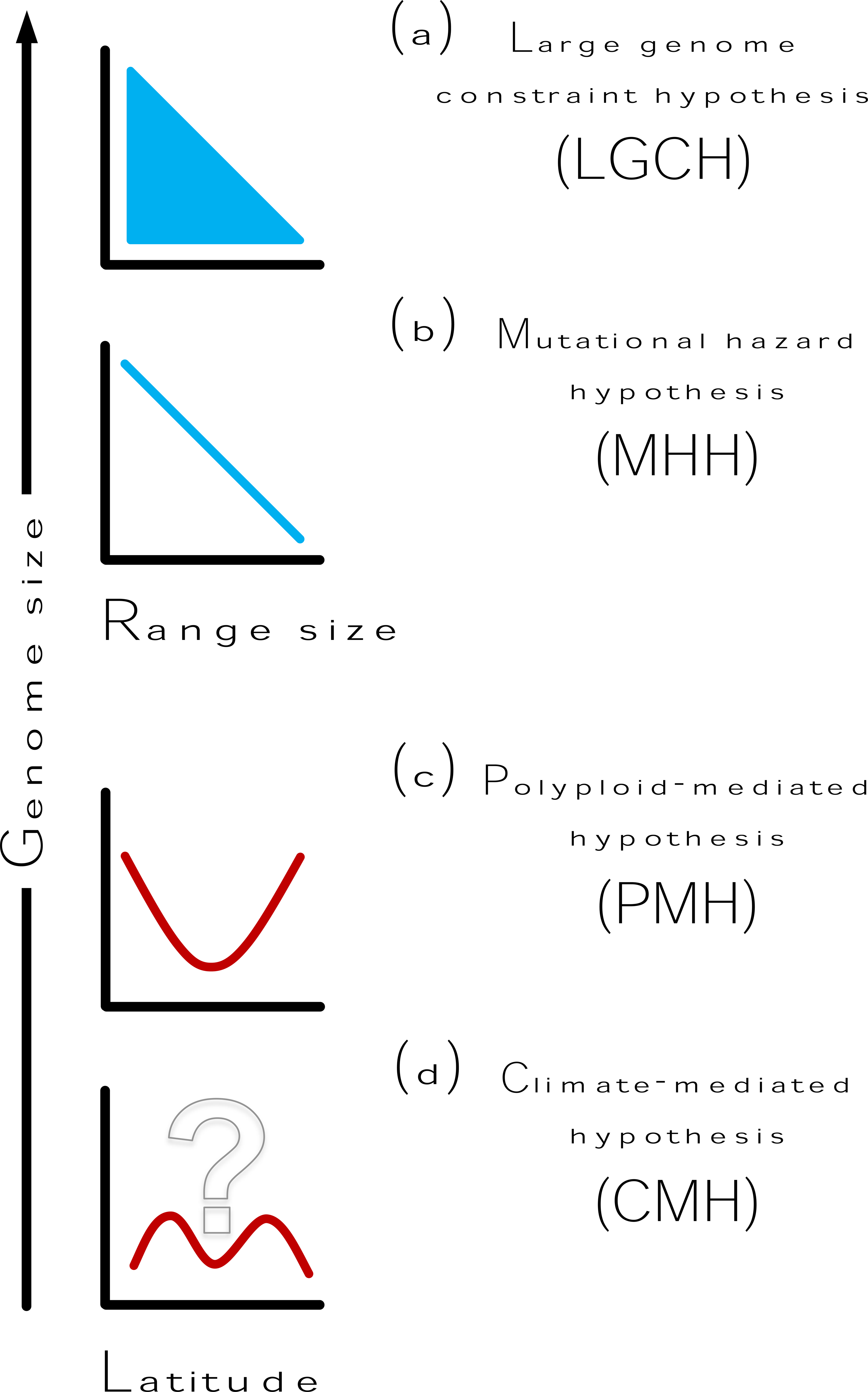
Expected associations between genome size and (a, b) range size and (c, d) latitude based on four hypotheses outlined in the Introduction. The question mark in (d) indicates uncertainty about the potential shape of the curve. Given this uncertainty, we present a curve that could possibly result from the effects of temperature.

Polyploidization is another major contributor to plant genome size evolution (Wendel, 2000; Leitch & Leitch, 2008; Soltis *et al*., 2015; Guignard *et al*., 2016; Šmarda *et al*., 2019), which, in newly formed polyploids (neopolyploids), leads to multiplication of the genome size and chromosome number (Mandáková & Lysák, 2018). However, over time, polyploids undergo post-polyploid diploidization that includes chromosome fusions and genome downsizing (Mandáková & Lysák, 2018), thereby blurring the clear correlation between genome size and chromosome number (Choi *et al*., 2020; Roddy *et al*., 2020). Because not all of the duplicated portion of the genome is eliminated during the post-polyploid diploidization (Bowers *et al*., 2003; Paterson *et al*., 2004; Wang *et al*., 2015), repeated polyploidization-diploidization cycles may lead to a gradual increase in genome size over time, especially in regions where polyploids originate more frequently. The proportion of neopolyploids at different latitudes across the globe shows a U-shaped pattern, being low in the tropics and increasing polewards (Rice *et al*., 2019). The latitudinal U-shape in the proportion of neopolyploids is likely a consequence of the similarly U-shaped distribution of the mechanisms underlying polyploid origin, for example, through the increased rate of formation of unreduced gametes at low temperatures (Ramsey & Schemske, 1998; Mason & Pires, 2015). As the relative positions of continents have remained similar over millions of years, latitudinal gradients in the rate of the repeated polyploidization-diploidization cycles (Wendel, 2015; Wendel *et al*., 2016; Clark & Donoghue, 2017) should persist over geological time scales and a U-shaped latitudinal distribution of genome size would gradually emerge in this scenario (Fig. 1c; polyploid-mediated hypothesis: PMH).

Latitudinal gradients encompass climatic and other environmental variables that could also be important factors contributing to genome size variation. These factors include temperature, precipitation, aridity, seasonality, ultraviolet-B radiation (UV-B), and length of the growing season (e.g., Bennett, 1976; Bennett *et al*., 1982; Grime & Mowforth, 1982; Rayburn & Auger, 1990; MacGillivray & Grime, 1995; Bottini *et al*., 2000; Knight & Ackerly, 2002; Grotkopp *et al*., 2004; Dušková *et al*., 2010; Díez *et al.,* 2013; Kang *et al*., 2014; Du *et al*., 2017; Bilinski *et al*., 2018; Souza *et al*., 2019; Becher *et al*., 2021; Cacho *et al*., 2021; Greimler *et al*., 2022; Sklenář *et al*., 2022). Studies of climatically-mediated (latitudinal or altitudinal) genome size distributions have found positive, negative, mixed, or quadratic responses of genome size to climatic gradients (reviewed in Cacho *et al*., 2021), which may be explained by their narrow geographic and taxonomic scopes (Knight & Ackerly, 2002; Greilhuber & Leitch, 2013). Nevertheless, one pattern that often emerges from these studies is the exclusion of the largest genomes from both ends of the climatic spectrum. This may arise from the complex ways in which the biophysical constraints imposed by genome size (e.g., setting the minimum cell size and duration of mitosis and meiosis) may impact many aspects of a plant’s biology, such as the timing of growth and physiological factors such as water and nutrient use efficiency, and hence influence where plants grow. For example, in the case of temperature, large genomes might be predicted to be excluded from areas with both the lowest and highest mean temperature (underpinned, in part, by the impact of genome size on the rate of cell division) (Fig. 1d; *climate-mediated hypothesis*: CMH).

Here, we test the following hypotheses (Fig. 1): 1) the *large-genome constraint hypothesis* (LGCH), which predicts species which occupy large geographical ranges cannot have large genomes; 2) the *mutational-hazard hypothesis* (MHH), which predicts that genome size decreases with increasing geographic range size; 3) the *polyploid-mediated hypothesis* (PMH), which predicts an increase in genome size from the equator to the poles; and 4) the *climate-mediated hypothesis* (CMH), which predicts the exclusion of large genomes from both ends of the climatic spectrum. We achieve this by combining the largest dataset compiled to date for angiosperm genome size (16,017 species) with newly-available data on the global distribution of angiosperms from the WCVP, and mapping the global distribution of angiosperm genome size.

## Material and Methods

### Taxonomic framework and geographic distribution

The angiosperm species nomenclature considered in this study follows the World Checklist of Vascular Plants (WCVP; Govaerts *et al*., 2021). We provide details of the accepted names, pertinent synonyms, and authorities for sampled taxa, as well as their WCVP ‘plant_name_id’ and distribution ranges based on LevelD3 Continental and Regional Codes (i.e., botanical countries) established by the International Working Group on Taxonomic Databases for Plant Sciences (TDWGs hereafter; Brummitt *et al*., 2001) in Supporting Information Dataset S1. This dataset also includes new validly-published species yet to be included in the WCVP database (marked as “NA” in the column “POWO ID” in Dataset S1), their distribution ranges converted to TDWGs, and corresponding sources. In exceptional cases when the WCVP taxonomic framework differed from the Catalogue of Life (Roskov *et al.,* 2019), World Plants (Hassler, 2022), or other sources, and this difference was supported by different genome sizes, we adopted the framework congruent with the genome size data (Dataset S1). We discarded taxa that were imprecisely identified (e.g., those only determined at the generic level), cultivated species with unknown native distributions, and hybrids (with the exception of a few cases where hybrid taxa have been accepted as species in some floras).

### Distributional range size estimation

Distribution range sizes were calculated as the extent of occurrence (EOO) for each species based on the Global Biodiversity Information Facility (GBIF) distribution data. To obtain EOO estimates in square kilometers, we first cleaned the data for species occurrences from GBIF following Elliott *et al*. (2022). Then, we calculated EOO (Dataset S1) using the ‘eoo’ function in the R package RANGEMAP v.0.1.18 (Cobos *et al*., 2022), with the ‘polygons’ option set to ‘simple_wmap(“simplest”)’ to omit oceans from the calculations. In addition, as an alternative measure of range size, we calculated the number of occupied TDWGs flagged as native for each species (Dataset S1).

### Genome size compilation

We extracted genome size estimates from several sources, including (1) research papers published between 2012 and 2022 (or older studies that were absent from Release 8.0 of the Angiosperm DNA C-values Database) retrieved using ‘Web of Science’, ‘ResearchGate’ and ‘Google Scholar’ (9,515 taxa, 59.4 %); (2) the Angiosperm DNA C-values Database (5,973 taxa, 37.3 %; Release 8.0: December 2012, Bennett & Leitch, 2012; Release 9.0: April 2019, Leitch *et al*., 2019), and (3) unpublished genome size measurements from the Plant Biosystematics Research Group of Masaryk University and the Royal Botanic Gardens, Kew (529 taxa, 3.3 %). Three different criteria were applied in cases where genome sizes for the same species were reported independently by different authors. These comprised (i) selecting values measured by flow cytometry over those estimated with Feulgen densitometry, (ii) choosing estimates from more recent reports over older ones, and (iii) assessing the taxonomic expertise of the authors for the species studied (i.e., we preferentially selected estimates from authors with taxonomic expertise in the group of interest when possible). We chose the smaller genome size (and thus the smaller DNA ploidy level) in cases where genome size varied within a species, corresponding to different DNA ploidy levels. For multiple estimates presented for a species in the same publication, the genome size values were averaged. Finally, in cases where publications used nomenclature that conflicted with the WCVP and genome size values reflected this difference, we chose an alternative taxonomic framework (predominantly the Catalogue of Life) and listed the source in Dataset S1. Genome size estimations reported in pg were converted to Mbp using the equation 1 pg = 978 Mbp (Doležel *et al.,* 2003).The genome size per TDWG was calculated as the average of the reported genome sizes for all taxa occurring in each region, which were log_10_-transformed (Dataset S2).

### Chromosome number compilation

Chromosome numbers were extracted (in order of preference) from: (i) the same publications as the genome size data when both estimates were reported together; (ii) the Chromosome Counts Database (CCDB: Rice *et al.,* 2015); and (iii) publications reporting only chromosome number (Dataset S1). We first ensured the estimations were not pseudo-replicated and then we selected the most prevalent number for a species. We report the median value for a species when it was not possible to discern the prevailing chromosome number (e.g., in cases of aneuploidy). When chromosome numbers varied based on differing ploidy levels within a species, we compared the ploidy levels and chromosome numbers of other congeners to aid in selecting the chromosome number corresponding to the reported genome size of that species. Finally, we calculated the mean chromosome size of a species by dividing the 2C genome size (in Mbp) by the diploid (2n) chromosome number. As mean chromosome size removes the correlation between genome size and chromosome number, we used it throughout the study as a correction for neopolyploidy (i.e., polyploids still recognizable cytologically rather than those with polyploidy in their ancestry recognizable only through DNA sequence analysis).

### Polyploid distributions

We extracted inferred ploidy-level data from Rice *et al*. (2019: https://figshare.com/collections/The_Global_Biogeography_of_Polyploid_Plants/4306004). Duplicate records and species that are not accepted in the WCVP were omitted from the dataset. We linked the remaining species to their geographic distribution based on TDWGs, as specified by the WCVP. We used the ploidy-level inferences to calculate the proportion of polyploids per TDWG (Dataset S2).

### Phylogenetic tree used in tests of MHH and LGCH

We used one hundred species-level trees of all angiosperms comprising all 329,798 species recognized by version 6 of the World Checklist of Vascular Plants (Forest, 2023) pruned to species in our dataset.

### Growth form classification

A relationship between genome size and growth form has been suggested by many authors (e.g., Bennett, 1971; 1987; Beaulieu *et al.,* 2008; Francis *et al.,* 2008, Veselý *et al.,* 2012; 2013). To control for this effect, all taxa were classified according to four plant growth forms (Dataset S1): (i) annuals (= therophytes; 12 % of species in the dataset), (ii) geophytes (11 %), (iii) non-geophytes (perennial herbs = hemicryptophytes + parasites + hydrophytes + epiphytes; 47 %), (iv) woody plants (= chamaephyte + phanerophytes; 30 %), using standard floras or The World Checklist of Selected Plant Families (WCSP, 2017). For each TDWG, we calculated the percentage of species belonging to the four growth forms (Dataset S2).

### Latitude estimations

We assigned a latitude to each TDWG (Dataset S2) using their geographic centroids, determined using ArcGIS v.10 (Environmental Systems Research Institute, 2014). The latitude associated with each species (Dataset S1) was then calculated as a mean of latitudinal centroids of all the TDWGs occupied by a given species.

### Climatic variables

We extracted 25 bioclimatic variables from the CHELSA database (Karger *et al*., 2017; https://chelsa-climate.org/bioclim/; Karger *et al*., s.a.), three ultraviolet-B-related variables from Beckmann *et al*. (2014; UVB1 = Annual Mean UV-B, UVB3 = Mean UV-B of Highest Month, and UVB5 = Sum of Monthly Mean UV-B during Highest Quarter), and the Global-Aridity Index (Global-Aridity_ET0; Trabucco & Zomer, 2018) at 30 arc-second resolution (∼1km). We then calculated the mean of each variable per TDWG region (Dataset S2) with QGIS v.3.14 “pi” (QGIS Development Team, 2022). Collinearity was then assessed by calculating Pearson correlation coefficients among all pairs of the 29 variables. Correlated variables (Pearson correlation coefficient > 0.7) were assembled into six groups (Fig. S1, Table S1). To select a single variable from the six groups for further analyses, we used each variable as a predictor of 2C genome size in a polynomial regression and selected those with the best explanatory power within their groups. To select an appropriate order of the polynomials for the regression, we used the cost function combined with a visual inspection of the bivariate plots of each variable and 2C genome size. We omitted GDD0 (Growing degree days heat sum above 0°C) and Aridity index from further consideration because both explained very little variation in the regression models (R_2ad_*_j_* = -0.002 and 0.001, respectively). Thus, the variables selected for further analyses (Table S1) were GST (Growing Season mean Temperature), BIO2 (mean diurnal air temperature range), BIO13 (precipitation of the wettest month), and BIO15 (precipitation seasonality).

Even if variables are collinear, the essence of their influence on genome size may differ (e.g., UV-B-caused deletion bias vs. temperature-affected cell size). Therefore, we performed additional analyzes with selected variables that did not pass the above-mentioned filtering steps (GSL – length of the growing season, UVB1 – mean annual UVB, BIO11 – Daily mean air temperatures of the coldest quarter), if they had biological relevance or their effect on genome size had already been hypothesized.

### Statistical analyses

We applied a series of linear regressions to test our four hypotheses (Fig. 1). The LGCH and MHH were modeled with genome size as a function of range size, with both variables log-transformed (base 10) to account for the skew towards low values. We first performed ordinary least squares regression (OLS) using the function ‘lm’ implemented in base R, followed by phylogenetic generalized least square (PGLS) regression (Freckleton *et al*., 2002) with the R package PHYLOLM v.2.6.2 (Ho & Ané, 2014). In PHYLOLM, we used the weighted Akaike information criterion (AICw; Akaike, 1978; Wagenmakers & Farrel, 2004) to select between seven evolutionary explicit models of trait evolution: Brownian motion, Pagel’s lambda, kappa, and delta, two Ornstein-Uhlenbeck models with an ancestral state estimated at the root or having the stationary distribution at the root, and the early burst model. The best model was Pagel’s lambda with AICw = 1 (averaged across all 100 trees), which we used to optimize branch lengths based on the data (model = ‘lambda’) using maximum likelihood estimation. To examine whether the association between range size and genome size is dependent upon differences in genome size, we applied quantile regression analysis with nineteen different quantiles (from 0.05 to 0.95 at 0.05 intervals) using function ‘rq’ in the R package QUANTREG v.5.93 (Koenker *et al.,* 2022). To the best of our knowledge, a tool has yet to be developed that is capable of performing quantile regression while correcting for evolutionary relationships among taxa. To circumvent this problem, we followed the multistep approach of Jovani *et al*. (2016), employing R packages CAPER v.1.0.1 (Orme, 2013) and QUANTREG v.5.93 (Koenker *et al.,* 2022).

To examine how genome size is associated with latitude (testing the PMH and CMH hypotheses), we specified genome size (log-10 transformed) as the response variable and latitude as the predictor variable in an OLS regression model. We used the cost function and the visual inspection of the bivariate plot of genome size and latitude to select the order of the polynomial fit and found that the best model was the third-degree polynomial (log_10_(Genome size)∼latitude+latitude_2_+latitude_3_). We also performed a multiple linear regression (MLR) that included the selected bioclimatic variables (i.e., GST, BIO2, BIO13, BIO15 - see above) as predictors to evaluate the potential effects of climatic factors on the distribution of genome size across latitude. In this MLR, we specified interaction terms among all predictor variables and conducted a backward stepwise model selection based on AIC values using the “step” function in base R. Based on the AICs from the backward selection process, the best model included only GST as a single predictor of 2C genome size (log_10_(Genome size)∼GST+GST_2_). In all MLRs with polynomials, we fitted orthogonal polynomials using the “poly” function in base R, but the “raw” parameter was set to “TRUE” to obtain parameter estimates corresponding to response variable units. Each TDWG was weighted in the regression analyses to account for the total number of species reported to occur in the region and the percentage of these species for which we have genome size or polyploid data. The weight was then calculated as the ratio of the number of species for which we have genome size data (or the proportion of polyploids) and the number of all species in the TDWG (Dataset S2). To evaluate causal relationships between the effects of GST and percentage of growth forms on mean genome size across TDWGs, we employed a path analysis approach using the R package LAVAAN v.4.2.3 (Rosseel, 2012).

## Results

### Sampling bias

We compiled the largest genome size dataset to date, encompassing >5% of known angiosperm species (Dataset S1). Large datasets of phylogenetic representation and traits, including genome size data, are latitudinally biased, with northern latitudes being more thoroughly sampled (Vasconcelos, 2022). To check how this may have affected our data, we compared the across-TDWG latitudinal distribution of range sizes of all angiosperms in the WCVP to that of the taxa in our genome size dataset. Both datasets show an increase in range size from south to north (Fig. S2).

### Genome size and range size (LGCH, MHH)

Genome size and range size exhibit a triangular relationship (Fig. 2a), indicating that species with small ranges can have any genome size, while species with large ranges only have small genomes (i.e., species with large genomes do not have large range sizes). The OLS regression model based on log-transformed data (Table 1) revealed a significant decrease in genome size with increasing range size (Fig. 2b). The slope from the PGLS analysis, although still significantly negative (*b* = -0.007, *P* = 1.31e-06), was flatter than that from the OLS (*b* = - 0.039, *P* < 2e-16), due to a strong phylogenetic signal (Pagel’s λ = 0.916) in the genome size/range size relationship (Table 1, Fig. S3a). Both ordinary (Fig. 2c, Table S2) and phylogenetic (Fig. S3b, Table S3) quantile regressions showed more negative slopes for higher quantiles of genome size, indicating that the relationship between genome size and geographical range size is genome size dependent - becoming increasingly negative as genome size increases; in accordance with the triangular relationship. Although the slopes started decreasing at the genome size quantile 0.5 for the ordinary quantile regression (Fig. 2c), in the phylogenetic quantile regression, the slope decreased continuously with increasing quantiles (Fig. S3b). When we used the number of occupied TDWGs as a measure of range size (instead of the EOO), we observed very similar results (Fig. S4, S5, Tables S4, S5), suggesting that, at least for our dataset, TDWG counts provide a reasonable proxy for range size.

**Fig. 2.**
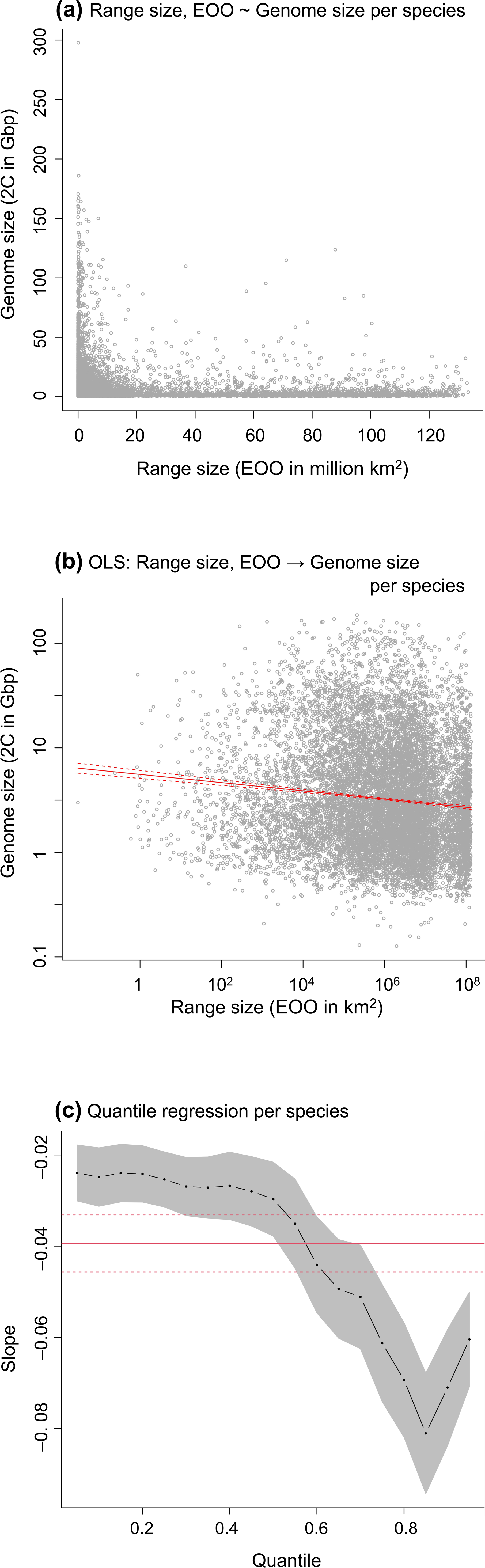
Associations between genome and range size per species (a, b, c). The association of the raw data between genome and range size is shown in (a), whereas both variables are log-transformed in the other two plots (b, c). The slope estimates from the quantile regression, including 95% confidence intervals (dark grey), are indicated in (c). The solid red line in (b) indicates the fit of the ordinary least squares (OLS) regressions, while the solid red line in (c) indicates the slope value from the OLS analysis. Dashed red lines (in b, c) represent 95% confidence intervals.

**Table 1:** Results of OLS and PGLS regressions of 2C genome size on range size. Table 1: Results of ordinary least squares (OLS) and phylogenetic generalised least squares (PGLS) regression of 2C genome size on range size. bi - regression estimates of model terms; 95%CI - lower and upper 95% confidence intervals of the regression estimates; R2adj - R squared adjusted indicating explained variance. The OLS analysis was perfromed with 12,137 species. The PGLS analysis was perfromed with 12,123 species. The PGLS was performed repeatedly with one hundred different trees (see Methods). Therefore, the values for PGLS are averages across these one hundred regressions.

| Table 1: Results of OLS and PGLS regressions of 2C genome size on range size |  |  |  |  |  |
| --- | --- | --- | --- | --- | --- |
| | OLS model: $\log_{10}(\text{2C genome size}) \sim \log_{10}(\text{Range size})$ | | | | |
| Model term | $b_i$ | 95%CI | t | P | $R^2_{adj}$ |
| Intercept | 3.746 | <3.708, 3.784> | 191.71 | <2E-16 | 0.012 |
| $\log_{10}(\text{Range size})$ | -0.039 | <-0.046, -0.033> | -12.24 | <2E-16 | |
| | PGLS model: $\log_{10}(\text{2C genome size}) \sim \log_{10}(\text{Range size})$ | | | | |
| Model term | $b_i$ | 95%CI | P | lambda | $R^2_{adj}$ |
| Intercept | 3.583 | <3.5782, 3.5887<br>> | <2E-16 | 0.916 | 0.002 |
| $\log_{10}(\text{Range size})$ | -0.007 | <-0.008, -0.0066<br>> | 1.31E-06 | | |
| Table 1: Results of ordinary least squares (OLS) and phylogenetic generalised least squares (PGLS) regression of 2C genome size on range size. $b_i$ - regression estimates of model terms; 95%CI - lower and upper 95% confidence intervals of the regression estimates; $R^2_{adj}$ - R squared adjusted indicating explained variance. The OLS analysis was performed with 12,137 species. The PGLS analysis was performed with 12,123 species. The PGLS was performed repeatedly with one hundred different trees (see Methods). Therefore, the values for PGLS are averages across these one hundred regressions. |  |  |  |  |  |

We also obtained very similar results when we controlled for the effect of neopolyploidy by performing the across-species analyses using mean chromosome size (2C/2n) instead of 2C genome size (Fig. S6, Tables S6-S8). For analyses with number of occupied TDWGs as a measure of range size, see Fig. S7, Tables S9-S11). However, the decrease in mean chromosome size with increasing range size was steeper than that of 2C genome size in both OLS and PGLS (Table 1, Table S6).

### Genome size, neopolyploidy, latitude, and climate

Overall, the smallest genomes occur in the tropics, and their size increases towards the poles. However, in the northern hemisphere, genome size decreases again from the temperate to the arctic regions. The global distribution of genome size averaged per TDWG is shown on the map in Fig. 3a. The genome size distribution maps of the two most species-rich eudicot (Asteraceae, Fabaceae) and monocot (Orchidaceae, Poaceae) families are shown in Fig. S8. Their genome size distribution resembles the overall trend in angiosperms. When the 2C genome size is plotted against the latitudinal centroids of TDWGs, the S-shaped pattern becomes evident (Fig. 3a). In the 3rd-order polynomial regression, latitude alone explained 40.12 % of the variation in 2C genome size (Table 2). The proportion of neopolyploid species displayed a U-shaped distribution with the smallest values in the tropics and a continuous increase in the proportion of polyploids towards the poles (Fig. 3b; Table 2).

**Fig. 3.**
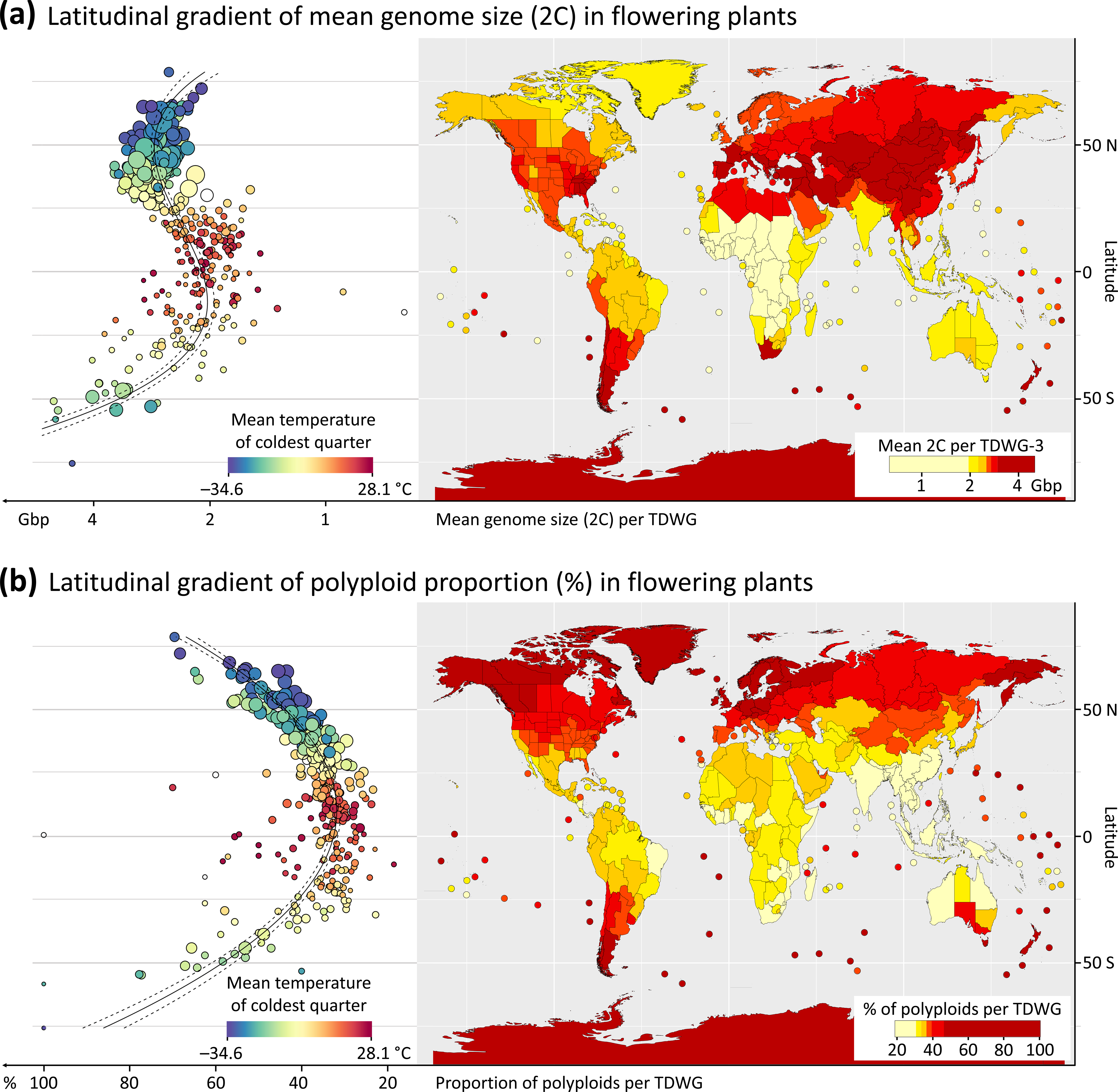
The global distribution of mean genome size (a) and polyploid proportion (b) in flowering plants. Mean genome size (2C) and the proportion of polyploids were calculated per TDWG Level-3 region. The two plots on the left side show (a) the distribution of genome size and (b) the proportion of polyploids across latitude. Dark red and dark blue indicate TDWG regions with the highest and lowest temperatures in the coldest quarter, respectively (BIO11 from CHELSA). The solid line in the plot indicates the mean from the regression fit. Dashed lines indicate 95% confidence intervals. The size of points in the plots indicates the weights used in the regression analysis. The weight was calculated as the ratio of the number of species for which we have genome size data (or the proportion of polyploids) to the number of all species in the TDWG. The maps on the right side show the distribution of (a) mean genome size and (b) polyploid proportion, with dark red and light yellow TDWG regions indicating areas with relatively high and low values for each variable, respectively.

**Table 2:**
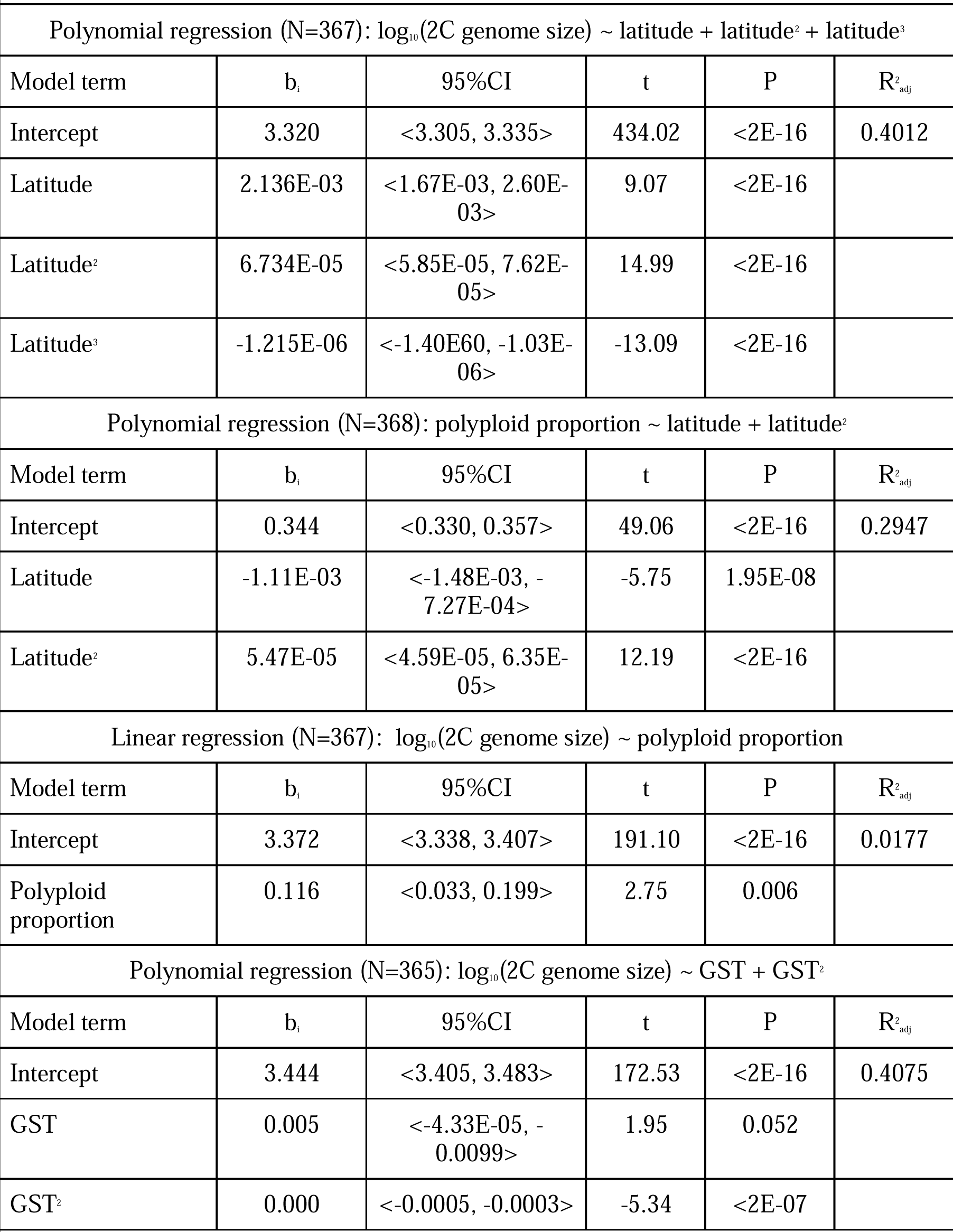
Results of linear and polynomial regressions of 2C genome size and polyploid proportion on various predictors. Table 2: N - number of TDWGs included in the analysis; bi - regression estimates of model terms; 95%CI - lower and upper limits of 95% confidence intervals of the regression estimates; R^2^_adj_ - R squared adjusted indicating explained variance. GST is the mean temperature of the growing season. In the case of polynomial regressions, we fitted orthogonal polynomials using the “poly” function in base R, but the parameter “raw” was set to “TRUE” to obtain parameter estimates corresponding to
response variables.

| **Table 2: Results of linear and polynomial regressions of 2C genome size and polyploid proportion on various predictors.** |  |  |  |  |  |
| --- | --- | --- | --- | --- | --- |
| Polynomial regression (N=367): $\log_{10}(\text{2C genome size}) \sim \text{latitude} + \text{latitude}^2 + \text{latitude}^3$ | | | | | |
| Model term | $b_i$ | 95%CI | t | P | $R^2_{\text{adj}}$ |
| Intercept | 3.320 | <3.305, 3.335> | 434.02 | <2E-16 | 0.4012 |
| Latitude | 2.136E-03 | <1.67E-03, 2.60E-03> | 9.07 | <2E-16 |  |
| Latitude<sup>2</sup> | 6.734E-05 | <5.85E-05, 7.62E-05> | 14.99 | <2E-16 |  |
| Latitude<sup>3</sup> | -1.215E-06 | <-1.40E-06, -1.03E-06> | -13.09 | <2E-16 |  |
| Polynomial regression (N=368): polyploid proportion $\sim \text{latitude} + \text{latitude}^2$ | | | | | |
| Model term | $b_i$ | 95%CI | t | P | $R^2_{\text{adj}}$ |
| Intercept | 0.344 | <0.330, 0.357> | 49.06 | <2E-16 | 0.2947 |
| Latitude | -1.11E-03 | <-1.48E-03, -7.27E-04> | -5.75 | 1.95E-08 |  |
| Latitude<sup>2</sup> | 5.47E-05 | <4.59E-05, 6.35E-05> | 12.19 | <2E-16 |  |
| Linear regression (N=367): $\log_{10}(\text{2C genome size}) \sim \text{polyploid proportion}$ | | | | | |
| Model term | $b_i$ | 95%CI | t | P | $R^2_{\text{adj}}$ |
| Intercept | 3.372 | <3.338, 3.407> | 191.10 | <2E-16 | 0.0177 |
| Polyploid proportion | 0.116 | <0.033, 0.199> | 2.75 | 0.006 |  |
| Polynomial regression (N=365): $\log_{10}(\text{2C genome size}) \sim \text{GST} + \text{GST}^2$ | | | | | |
| Model term | $b_i$ | 95%CI | t | P | $R^2_{\text{adj}}$ |
| Intercept | 3.444 | <3.405, 3.483> | 172.53 | <2E-16 | 0.4075 |
| GST | 0.005 | <-4.33E-05, -0.0099> | 1.95 | 0.052 |  |
| GST<sup>2</sup> | 0.000 | <-0.0005, -0.0003> | -5.34 | <2E-07 |  |
| Table 2: N - number of TDWGs included in the analysis; $b_i$ - regression estimates of model terms; 95%CI - lower and upper limits of 95% confidence intervals of the regression estimates; $R^2_{\text{adj}}$ - R squared adjusted indicating explained variance. GST is the mean temperature of the growing season. In the case of polynomial regressions, we fitted orthogonal polynomials using the "poly" function in | | | | | |

Genome size and the proportion of polyploid species exhibited very different latitudinal distributions (Fig. 3), with the proportion of polyploid species explaining only 1.77 % of the variation in 2C genome size (Table 2).

When we controlled for neopolyploidy by analyzing mean chromosome size across TDWGs, the S-shape latitudinal trend remained broadly unchanged (Fig. S9). The S-shaped latitudinal trend in genome size was robust to longitude, as the same pattern was recovered when the data were separately analyzed for the New and Old Worlds (Fig. S10).

To assess which climatic parameters might be associated with the observed latitudinal trend in 2C genome size, we tested 29 climatic variables, but only GST (mean temperature of the growing season) was used in the final regression model based on backward selection (see Methods for details). The best-fitting model was a quadratic polynomial regression of 2C genome size on the GST (Table 2). The quadratic term had a negative coefficient, indicating that genomes are smaller in TDWGs with high or low temperatures and larger for intermediate temperatures (Table 2; compare with the graph in Fig. 2a). The GST explained 40.75 % of the variance in 2C genome size which is all the variance explained by latitude (40.12 %; Table 2). If BIO11, which falls below 0°C in the northern hemisphere, is added into the model, the explained variance increases to 46.35 % (Table S12), highlighting the importance of freezing temperatures. Furthermore, if the MHH is combined with CMH by adding the range size to the model with GST, the explained variance increases to 46.14 % (Table S12).

We also tested whether smaller genomes are linked to shorter growing seasons. Our regression analysis showed that as the genome gets larger, the growing season (GSL) gets shorter (P=0.0004; Table S12). When analyzed only for TDWGs with latitudinal centroids of at least 48.93° (the threshold at which genomes start decreasing northward), genome size decreases with a shortening of the growing season, but the relationship is not significant (P=0.481; Table S12). UVB1 (mean annual UVB) explained 34.6 % of the variation in mean genome size across TDWGs (Table S12).

Due to genome size variations among different plant growth forms (Bennett, 1987; Beaulieu *et al*., 2008; Veselý *et al*., 2013), and the presence of latitudinal trends in growth form proportions (Taylor *et al*., 2023; Fig. S11 here), we investigated whether the observed S-shape (Fig. 3a) might be attributed to differences in the percentages of different growth forms within TDWGs with increasing latitudes. Annuals, geophytes, and non-geophyte herbs all exhibited the S-shape in mean genome size, varying only in magnitude (Fig. 4). Woody plants, however, had slightly larger genomes in the tropics compared to temperate or arctic regions (Fig. 4). These growth form patterns remained consistent across both species (Fig. 4a) and TDWG means (Fig. 4b-e). As sole predictor, the percentage of growth forms explained from 2% of genome size variance (in annuals) to 21% (in non-geophytes) (Table S13). However, when growth form percentage was added to the model with GST, the effects of non-geophytes and annuals became insignificant, with geophytes and woody plants contributing only 3.4% and 1.4% additional explained variance, respectively (Table S14). This significant drop in the explanatory power of growth forms suggests that GST directly influences both growth form percentages and mean genome size within TDWGs. This was confirmed through path analysis, which revealed that while GST strongly impacts genome size and the percentages of non-geophytes and woody plants, growth forms have minimal or negligible effects on the distribution of genome sizes across the globe (Fig. S12).

**Fig. 4.**
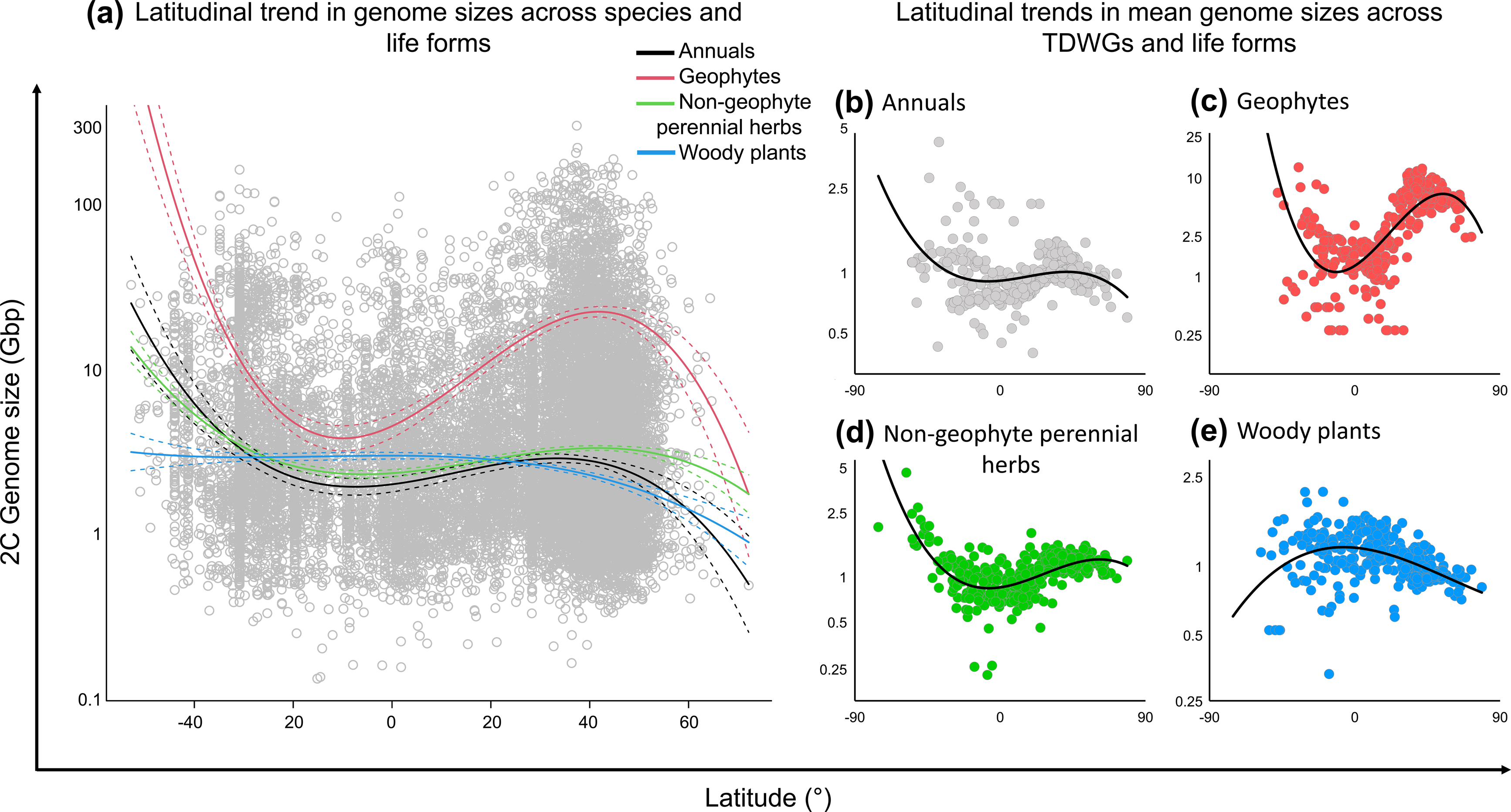
The association of genome size (2C; Gbp) and latitude across four growth forms groupings. Plot (a) is based on species genome sizes (grey circles), whereas the latter four plots (b, c, d, e) represent the mean genome size calculated per TDWG for a given growth form. All results are based on polynomial regressions of the 3_rd_ order, where solid lines represent the model estimates. The dashed lines in (a) show the 95% confidence intervals.

## Discussion

### Support for the LGCH, while not ruling out the MHH

We revealed a triangular relationship between range size and genome size, with a negative association between range size and genome size that is accentuated as genome sizes increase (Fig. 2b, 2c), supporting the LGCH (Fig. 1a). This relationship indicates that large-genomed species are restricted to occupying smaller ranges, which is likely due to the nucleotypic effects of their genomes hindering their dispersal distance and limiting their ecological niche (Knight & Ackerly, 2002; Beaulieu *et al*., 2007, 2008; Veselý *et al.,* 2012; Carta *et al.,* 2022; Bhadra *et al*., 2023). This places large-genomed species at a disadvantage compared to their smaller-genomed counterparts that have greater nucleotypic plasticity (Mayerson *et al.,* 2020; Bhadra *et al*., 2023) and may thus occupy both large and small ranges (Fig. 2a). It is notable that the most pronounced S-shape in the latitudinal distribution of genome size (see *Genome size decreases* […] *but not in the south* section below) is in geophytes (Fig. 4c), whose genomes are the largest among the analyzed growth forms (Fig. 4a). Although the triangular relationship we observed does not show support for the MHH, the LGCH does not necessarily rule out the MHH. Notably, the largest genomes are found in the southern hemisphere (Fig. 3a), where angiosperms in our dataset have the smallest ranges (Fig. S2) and could thus be most susceptible to genetic drift (Fig. 5). Genetic drift could facilitate genome growth in smaller-ranged species (as proposed in the MHH), which could further reduce the range size of large-genomed species (LGCH) and throw them into a deadly descending spiral toward extinction. This is supported by evidence showing that large-genomed species are at higher risk of extinction (Vinogradov, 2003; Soto Gomez *et al*., 2023 in this issue).

**Fig. 5.**
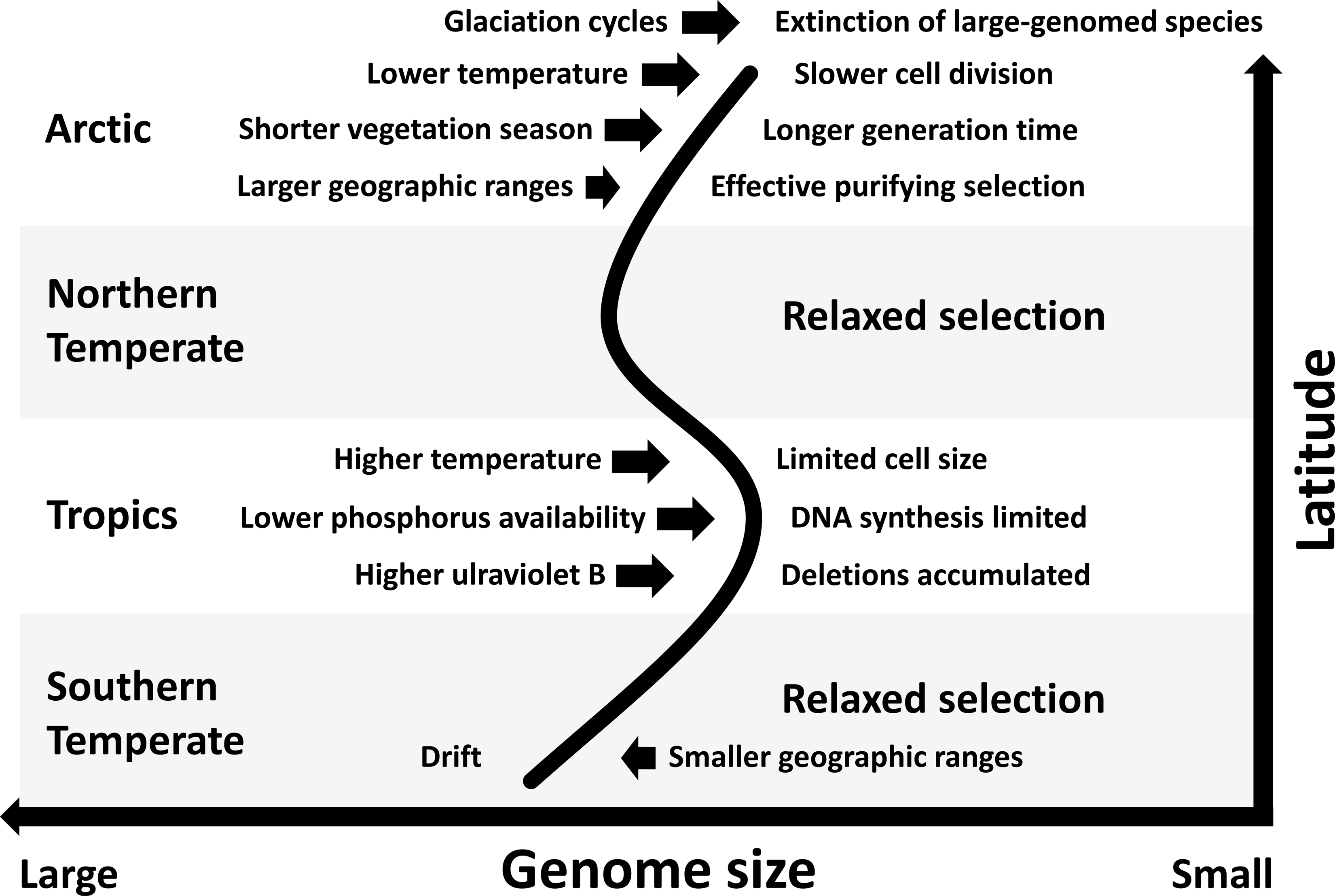
Proposed major factors (to the left of arrows) affecting physiological, anatomical, and molecular response (to the right of arrows), resulting in expansions/contractions of the genome and thus ultimately forming the global latitudinal trend in genome size (S-shaped curve). Relatively high genome sizes in the temperate regions could be the result of relaxed selective pressure, whereas various drivers might be constraining or pushing this trait in tropical and arctic regions. The proportion of polyploid species (low in the tropics and increasing toward the poles), which is not included in the figure, could also weakly contribute to the observed latitudinal trend in genome size (see Table 2).

### Small genomes in the tropics

The decrease in genome size from temperate to tropical regions across both hemispheres is consistent with previous studies focusing on genome size (or its proxies) in Poaceae (e.g., Avdulov, 1931; Bennett, 1976), Fabaceae (e.g., Stebbins, 1966; Bennett, 1976; Souza *et al*., 2019), Orchidaceae (e.g., Trávníček *et al*., 2019), Zygophyllaceae (e.g., Vidal-Russell *et al*., 2022) and at broader phylogenetic scales across angiosperms (Levin & Funderburg, 1979; Yu *et al.,* 2018). In our study, the environmental variables most correlated with latitude were temperature-related, and peaked in the tropics (Fig. 3a). In higher ambient temperatures, metazoan ectotherms, unicellular eukaryotes, and prokaryotes tend to have smaller cells (Atkinson *et al*., 2003; Hessen *et al*., 2013; Sabath *et al*., 2013), possibly because the maintenance of large cells becomes more difficult with increasing temperatures (Sabath *et al*., 2013). Our finding that small genomes are prevalent in the tropics might reflect this relationship, where it is advantageous to have smaller cells (and thus a smaller genome; Cavalier-Smith, 2005) in the tropics.

Alternatively, small genomes in low latitudes could be a result of the DNA-damaging effects of UV-B radiation (Bennett, 1976), which is generally highest in the tropics (Beckmann *et al*., 2014) and might result in selecting for smaller chromosomes that absorb less energy, therefore decreasing radiosensitivity (Sparrow *et al*., 1967). This idea is supported by recent findings showing that plants with holocentric chromosomes, which tolerate fragmentation (Zedek & Bureš, 2019), are less stressed (Zedek *et al.,* 2020; 2021) and more competitive (Zedek *et al.,* 2022) under higher UV-B doses. Moreover, homologous recombination used to repair UV-B-induced damage might increase rates of DNA deletion, thereby further promoting genome downsizing (Schubert & Vu, 2016). However, as UV-B radiation intensity (which explained 34.6 % of the variation) is strongly correlated with temperature (Fig. S1), the individual effects of these two factors on genome size in the tropics cannot be easily differentiated.

Finally, nutrient limitation might play a role in constraining the genome size of tropical plants, as many tropical soils are low in nutrients (especially phosphorus; Vitousek *et al*., 2010), and yet building and maintaining cells in plants with large genomes is expensive in terms of N and P. This may result in species with large genomes being less competitive in the nutrient-poor tropical soils, resulting in their exclusion from these environments (Leitch & Leitch, 2008; Šmarda *et al*., 2013, Guignard *et al*., 2016; Faizullah *et al*., 2021; Veleba *et al*., 2020).

### Genome size decreases from temperate regions towards the North pole, but not the South

Differences in genome size trends across latitudinal gradients in the northern versus southern hemisphere may be explained by the larger temperature gradient in the north, which could be partially associated with differences in the distribution of landmasses and major water bodies in the two hemispheres. Large areas of Eurasia and North America extend beyond 50°N and are surrounded by less water and more land masses than regions in the southern hemisphere, therefore experiencing weaker buffering effects from the ocean. If the distribution of genome sizes in plants followed a similar pattern to the distribution of polyploid species, genomes would be expected to be smaller near the equator and increase polewards. We find that this trend holds, but only up to a certain, presumably limiting, low temperature threshold, beyond which genome sizes decrease towards the high northern latitudes. In the southern hemisphere, this low-temperature threshold is probably not reached (Fig. 1a). The existence of such a latitudinal breakpoint in genome size was previously predicted (Bennett *et al.,* 1982). The main drivers of selection pressure against larger genomes in polar regions were predicted to be: (i) slower cell divisions mediated by lower temperatures (Francis & Barlow, 1988) and (ii) longer generation times mediated by lower temperatures and/or by shorter growing seasons (Bennett *et al*., 1982; Bennett, 1987). Indeed, temperature variables alone explain a relatively large proportion (up to ∼40 %) of the variation in the global distribution of genome sizes (Table 2, Table S1).

Several authors have hypothesized that the decrease in temperatures toward the poles can result in a higher production of unreduced gametes in plants (Belling, 1925; Sakamura & Stow, 1926; de Mol, 1928; Heilborn, 1930; Hagerup, 1932; Bretagnolle & Thompson, 1995; Mason & Pires, 2015; Kreiner *et al*., 2017). If this phenomenon explains the increase in the proportion of polyploidy from the equator to the poles (Fig. 3b; Rice *et al*., 2019), then the polyploid proportion should be significantly higher in the northern hemisphere, where temperatures reach lower values. However, neither our study nor that by Rice *et al*. (2019) found a difference in the proportion of polyploids between the southern and northern hemispheres (Fig. 3b), suggesting that unreduced gamete production might not be an important variable in explaining latitudinal variation in polyploidy (but see below).

The decrease in genome size in the northern hemisphere from temperate regions to the Arctic could also be related to glaciation cycles, as smaller-genomed species tend to occur in previously glaciated TDWGs (Fig. S13). During glacial migrations, species with large genomes might have been more prone to extinction because of their smaller range sizes, as suggested by the negative association between geographic range size and genome size (Fig. 2a). Similarly, repeated glaciation cycles could have led to the extinction of some (neo)polyploids whose genome sizes exceeded an upper selection limit, which could further explain why the proportion of polyploids is not higher in the northern than the southern hemisphere. In this case, the hypothesis relating the increased formation of unreduced gametes to low temperatures and its role in increasing the proportion of polyploids from tropical to polar regions would still be relevant. A further possibility explaining the decrease in genome size from the northern temperate to polar regions is that shorter growing seasons towards high latitudes might be important in selecting plants with smaller genomes, which have faster growth rates and can complete their growth cycles in less time (Knight *et al.,* 2005). Nevertheless, our results show that any effect of length of growing season in the Arctic on genome size, is likely to be minor (Table S12).

### Relatively large genomes in temperate regions

Temperate regions offer mild conditions between the extremes of the tropics and arctic regions discussed above. For instance, there are not very high nor low temperatures, lower doses of UV-B radiation than in the tropics, and the area was not as extensively glaciated as arctic regions. The temperate climate might thus relax selective pressures against larger genomes, thereby increasing the overall range and mean genome sizes of plants growing in temperate zones of both hemispheres (Fig. 4a).

### Latitudinal gradient in genome size is not underpinned by contrasting proportions of different growth forms in different regions

Although the proportion of growth forms, especially perennial herbs and woody plants, may have significantly contributed to the global distribution of polyploids (Rice *et al*., 2019), the impact of different growth forms on the global distribution of genome size appears weak and mostly mediated by temperature (Fig. S12, Table S14). The independence of global genome size distribution on growth forms is further supported by the observation that annuals, geophytes, and non-geophytes all exhibited the S-shape in mean genome size (Fig. 4). Woody plants showed a different pattern, but their genome size still decreased northward (Fig. 4). Woody angiosperms are seldom polyploid (Müntzing, 1936; Stebbins, 1940; Otto & Whitton, 2000; Zenil-Fergusson *et al*., 2017; Rice *et al*., 2019), which could explain why their genomes did not increase in temperate regions. Also, the absence of relationship between extinction risk and genome size in woody plants (Soto Gomez *et al*., 2023) could suggest that genome size dynamics operate differently in woody vs herbaceous species.

### Conclusions and future directions

Our study found support for the large genome constraint hypothesis in explaining the global distribution of genome sizes but could not rule out the mutation hazard hypothesis in also contributing to explaining the distribution patterns observed. In addition, we show a small effect of polyploidy and growth forms and a large effect of climate, especially temperature, on the distribution of genome size. Overall, our findings indicate that mainly purifying selection, genetic drift, relaxed selection, and environmental filtering influenced by climate are likely to have shaped the global distribution of angiosperm genomes sizes (Fig. 5). Further research should be directed at determining the relative contributions of long-term processes shaping the global distribution of genome sizes, such as glaciation cycles, UV-B-caused genome erosion, or polyploidization-rediploidization cycles. We also advocate more thorough investigation of links between environmental factors and genome size at finer regional or local scales. For instance, the use of vegetation plots combined with species Ellenberg indicator values would enable a more in-depth understanding of the complex interplay between genome size and both biotic (e.g., competition) and abiotic (e.g., altitude, temperature, soil reaction and moisture) factors in influencing a species habitat and niche and its resilience to environmental changes.

## Author contributions

PB, TLE, and FZ designed the study, performed the analyses, and drafted the first version of the manuscript. PB and FZ collected the genome size data. PV and PB prepared the datasets for analysis, and PV assigned growth forms and contributed to the analysis. FF prepared phylogenetic trees. JŠ performed flow cytometric measurement generating unpublished data used in this study. PŠ, FF, IJL, ENL, MSG, SP, and MJMB contributed to analyses, interpretation of the results, and the final form of the manuscript.

## Acknowledgments

We thank Rafaël Govaerts for providing data from the World Checklist of Vascular Plants. This work was financially supported by the Czech Science Foundation, grant no. GA20-15989S.

## Supporting Information

Additional Supporting Information may be found online in the Supporting Information section at the end of the article.

**Fig. S1** Pearson’s correlation coefficients (*r*) among 29 climatic variables assessed to be included in the multiple linear regression model explaining genome size variation along the global latitudinal gradient. Dark red and dark blue circles indicate high and low *r* values, respectively. Larger circles in the upper triangle represent stronger correlations between variables (both negative and positive), whereas the numbers in the lower triangle indicate the *r* values.

**Fig. S2** Global distribution of mean geographic range sizes for those species included in the genome size dataset (a) and for all species in the WCVP dataset (b) mapped per TDWG Level-3 region. The two plots on the left-hand side of the figure show the distribution of mean geographic range sizes across the global latitudinal gradient. Dark red shading in the maps on the right-hand side of the figure indicates relatively high mean range sizes of species included in each TDWG unit, whereas light yellows indicate TDWGs with species with relatively small range sizes.

**Fig. S3** Associations between genome and range size (as Extent of Occurrence, EOO) per species considering phylogenetic relationships. The solid red line in (a) indicates the fit of the phylogenetic generalized least squares regression (PGLS), while the red line in (b) indicates the slope value from the phylogenetic quantile regression analysis. The slope estimates from the phylogenetic quantile regression, including 95% confidence intervals (error bars), are indicated in (b). Dashed red lines represent 95% confidence intervals. Both genome and range size are transformed by log_10_ in (a) and (b).

**Fig. S4** Associations between genome and range size per species when the number of occupied TDWG regions (instead of Extent of Occurrence, EOO) is used as a measure of range size. The association of the raw data between genome and range size is shown in (a), whereas both variables are log-transformed in the other two plots (b, c). The slope estimates from the quantile regression, including 95% confidence intervals (dark grey), are indicated in (c). The solid red line in (b) indicates the fit of the ordinary least squares (OLS) regressions, while the solid red line in (c) indicates the slope value from the OLS analysis. Dashed red lines represent 95% confidence intervals.

**Fig. S5** Associations between genome and range size per species considering phylogenetic relationships when the number of occupied TDWG regions (instead of the Extent of Occurrence, EOO) is used as a measure of range size. The solid red line in (a) indicates the fit of the phylogenetic generalized least squares regression (PGLS), while the red line in (b) indicates the slope value from the phylogenetic quantile regression analysis. The slope estimates from the phylogenetic quantile regression, including 95% confidence intervals (error bars), are indicated in (b). Dashed red lines represent 95% confidence intervals. Both genome and range size are transformed by log10 in (a) and (b).

**Fig. S6** Associations between mean chromosome size and range size (as Extent of Occurrence, EOO) per species. The solid red lines in (a) and (c) indicate the fit of the ordinary least squares (OLS) and phylogenetic generalized least squares (PGLS) regressions, respectively. The solid black circles and the gray shading in (b) represent the slope estimates and the 95% confidence intervals across 19 quantiles, whereas the hollow circles and the error bars in (d) indicate slope estimates and the 95% confidence intervals of the phylogenetic quantile regression. The horizontal red line in (b) represents the slope estimate of the OLS regression, while the horizontal red line in (d) shows the slope estimate of the PGLS regression. Dotted red lines in all four plots indicate the 95% confidence intervals of the slope estimates.

**Fig. S7** Associations between mean chromosome size and range size (as the number of occupied TDWG regions) per species. The solid red lines in (a) and (c) indicate the fit of the ordinary least squares (OLS) and phylogenetic generalized least squares (PGLS) regressions, respectively. The solid black circles and the gray shading in (b) represent the slope estimates and the 95% confidence intervals across 19 quantiles, whereas the hollow circles and the error bars in (d) indicate slope estimates and the 95% confidence intervals of the phylogenetic quantile regression. The horizontal red line in (b) represents the slope estimate of the OLS regression, while the horizontal red line in (d) shows the slope estimate of the PGLS regression. Dotted red lines in all four plots indicate the 95% confidence intervals of the slope estimates.

**Fig. S8** Mean genome sizes (2C; Gbp) averaged per TDWG region for the two most species-rich monocot (a – Orchidaceae, b – Poaceae) and dicot (c – Asteraceae, d – Fabaceae) families. Dark red colors indicate relatively large mean genome sizes, whereas light yellow shades indicate TDWG regions with relatively small mean genome sizes.

**Fig. S9** The global distribution of mean chromosome size in flowering plants calculated per TDWG region. The plot on the left side shows the distribution of mean chromosome sizes across latitudes, with dark reds indicating TDWG regions with high temperatures in the coldest quarter (BIO11 from Bioclim) and dark blues showing regions with low temperatures. The size of points in the plots indicates the weights used in the regression analysis (see Methods for details). The map on the right side shows the distribution of mean chromosome sizes mapped according to each TDWG region, where dark reds indicate relatively high values.

**Fig. S10** Mean genome sizes (2C; Gbp) across the global latitudinal gradient for the Old World (a) and New World (b). Circles in both plots represent the genome size averaged per TDWG region.

**Fig. S11** Latitudinal distribution of the percentage of (a) nongeophyte, (b) annual, (c) geophyte, and (d) woody species in our genome size dataset (Dataset S2).

**Fig. S12** Path analysis of causal relationships among the effects of the growing season temperature (GST) and percentages of species of different growth forms on the mean genome size in TDWG regions: (a) nongeophytes, (b) annuals, (c) geophytes, and (d) woody species. The numbers indicate standardized regression coefficients from the path analyses. The arrows show the direction of the causal effects, their thickness indicates the relative effects, the fading indicates significance of the effect and the color indicates positive (red) or negative (blue) effect.

**Fig. S13** Mean genome sizes (2C; Gbp) across the global latitudinal gradient illustrating TDWG regions that were glaciated (blue) and non-glaciated (red) during the last glacial maximum (LGM) approximately 18,000 years before the present. We assessed the glaciation status of each TDWG region at the Last Glacial Maximum (LGM; ∼18,000 years BP) using past climatic reconstructions from Ehlers (2015). We considered TDWG regions to be ‘Glaciated’ if their centroids were covered by the ice sheets during the LGM (Dataset S2).

**Dataset S1** Dataset containing 16,017 angiosperm taxa, their genome sizes, chromosome numbers, chromosome sizes, geographic ranges, latitudinal centroids, and growth forms.

**Dataset S2** Dataset containing 369 TDWGs (Botanical countries), their geographic centroids, counts of all angiosperm taxa and counts of angiosperm taxa with genomic traits; mean values for genome size, chromosome size, range size; mean values for genome size in growth forms; proportion of polyploid taxa; glaciation status; growth form percentages in TDWG regions.

**Table S1** Bioclim variables as they explain the variance in 2C genome size across TDWG regions in the polynomial regression of a given order.

**Table S2** Results of quantile regression of 2C genome size on range size (EOO).

**Table S3** Results of phylogenetic quantile regression of 2C genome size on range size (EOO).

**Table S4** Results of quantile regression of genome size on range size (TDWGs).

**Table S5** Results of phylogenetic quantile regression of genome size on range size (TDWGs).

**Table S6** Results of OLS and PGLS regressions of mean chromosome size on range size (EOO).

**Table S7** Results of quantile regression of mean chromosome size on range size (EOO).

**Table S8** Results of phylogenetic quantile regression of mean chromosome size on range size (EOO).

**Table S9** Results of OLS and PGLS regressions of mean chromosome size on range size (TDWGs).

**Table S10** Results of quantile regression of mean chromosome size on range size (TDWGs).

**Table S11** Results of phylogenetic quantile regression of mean chromosome size on range size (TDWGs).

**Table S12** Additional regressions of 2C genome size on other biologically relevant variables.

**Table S13** Results of regressions of 2C genome size on percentage of growth forms in TDWGs.

**Table S14** Results of regressions of 2C genome size on additive effects of GST and percentage of growth forms in TDWGs.

