## Supporting Information for "The global distribution of angiosperm genome size is shaped by climate"

### New Phytologist Supporting Information

Article acceptance date: XX XXX XXXX

The Following Supporting Information is available for this article

**Fig. S1** Pearson's correlation coefficients (r) among 29 climatic variables assessed to be included in the multiple linear regression model explaining genome size variation along the global latitudinal gradient. Dark red and dark blue circles indicate high and low r values, respectively. Larger circles in the upper triangle represent stronger correlations between variables (both negative and positive), whereas the numbers in the lower triangle indicate the r values.

**Table S12** Additional regressions of 2C genome size on other biologically relevant variables.

**Table S13** Results of regressions of 2C genome size on percentage of growth forms in TDWGs.

**Table S14** Results of regressions of 2C genome size on additive effects of GST and percentage of growth forms in TDWGs.

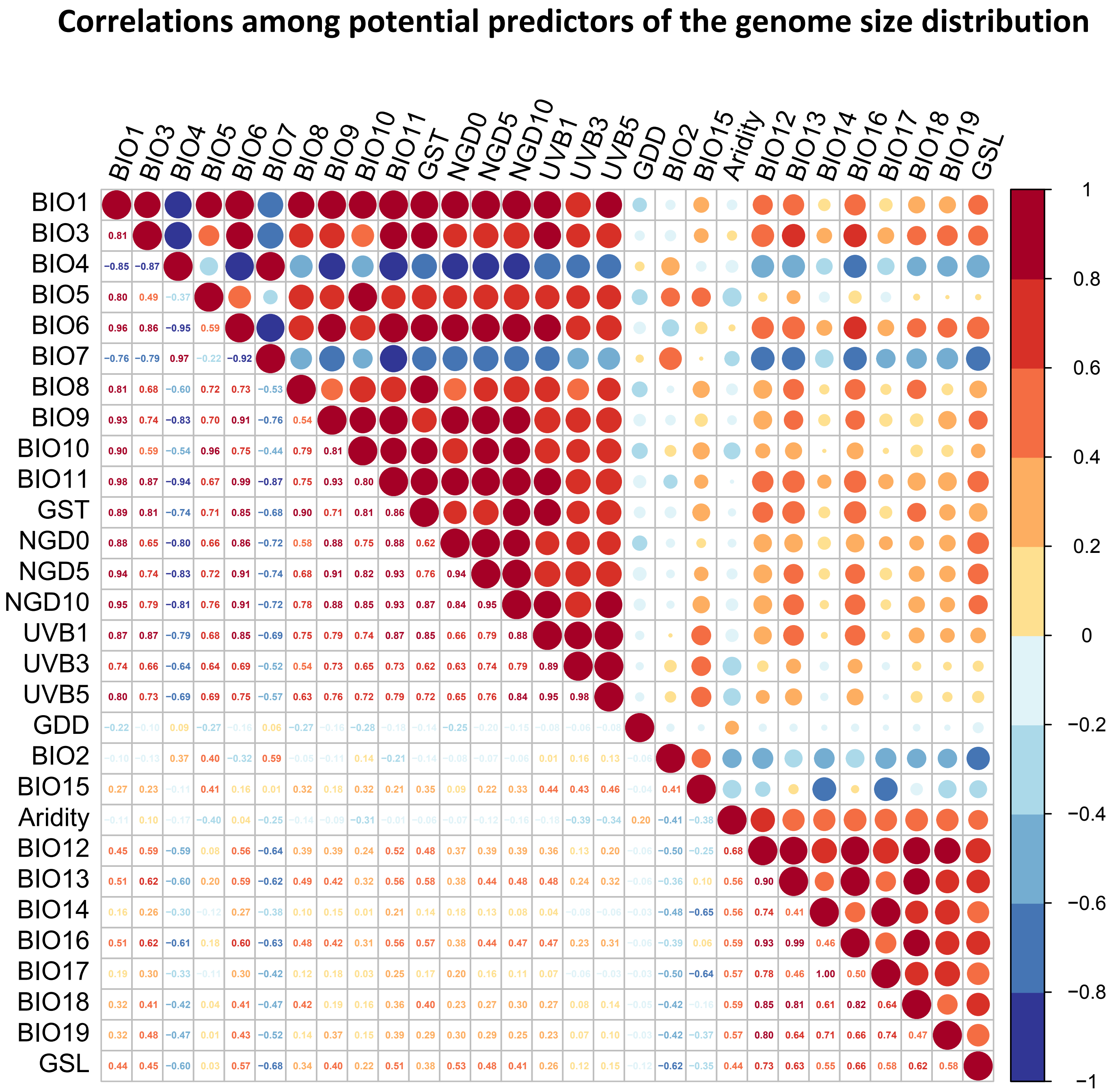


**Fig. S1** Pearson's correlation coefficients (*r*) among 29 climatic variables assessed to be included in the multiple linear regression model explaining genome size variation along the global latitudinal gradient. Dark red and dark blue circles indicate high and low *r* values, respectively. Larger circles in the upper triangle represent stronger correlations between variables (both negative and positive), whereas the numbers in the lower triangle indicate the *r* values.

#### Global distribution and latitudinal trend of geographic range sizes in flowering plants

**(a)** Gradient/distribution of mean geographic range size in flowering plants with genome size

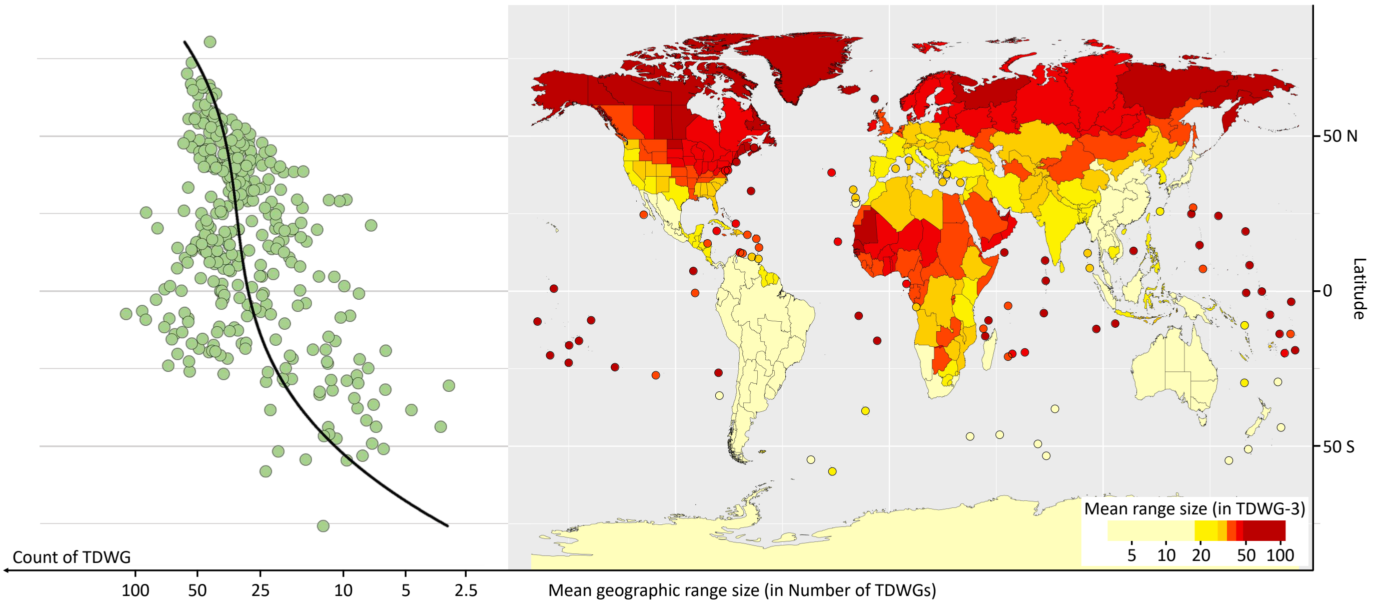


**(b)** Gradient/distribution of mean geographic range size across all flowering plants

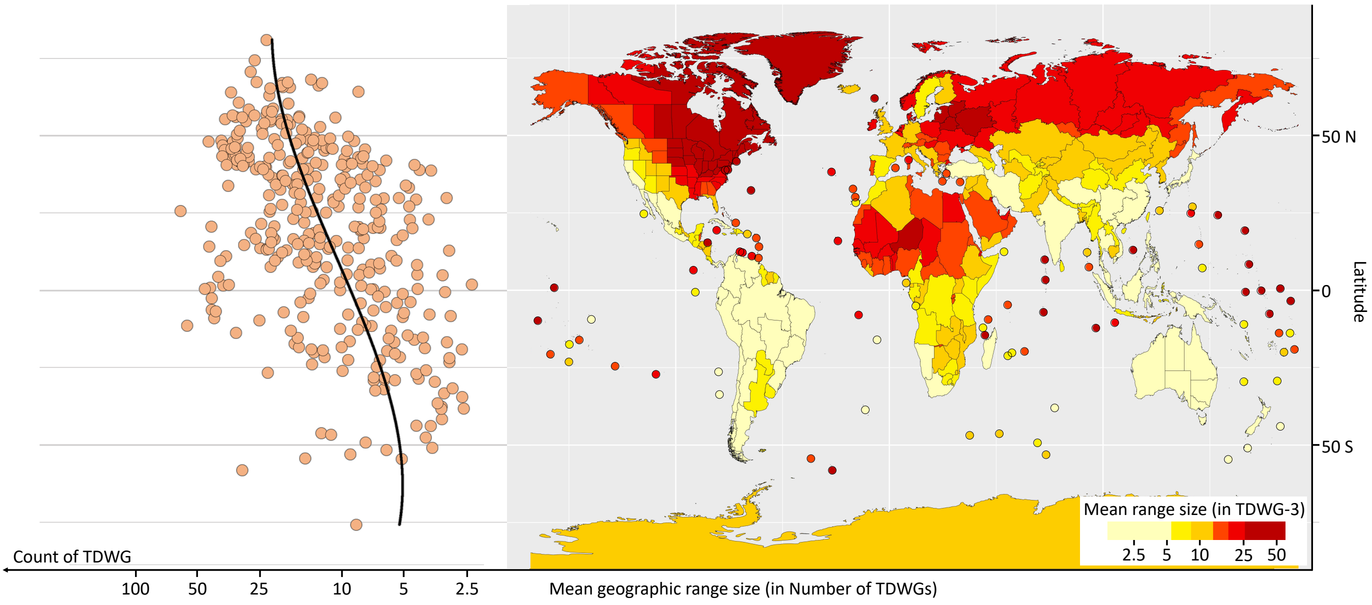


**Fig. S2** Global distribution of mean geographic range sizes for those species included in the genome size dataset (a) and for all species in the WCVP dataset (b) mapped per TDWG Level-3 region. The two plots on the left-hand side of the figure show the distribution of mean geographic range sizes across the global latitudinal gradient. Dark red shading in the maps on the right-hand side of the figure indicates relatively high mean range sizes of species included in each TDWG unit, whereas light yellows indicate TDWGs with species with relatively small range sizes.

### Associations between genome and range size (as Extent of Occurence, EOO) considering phylogenetic relationships

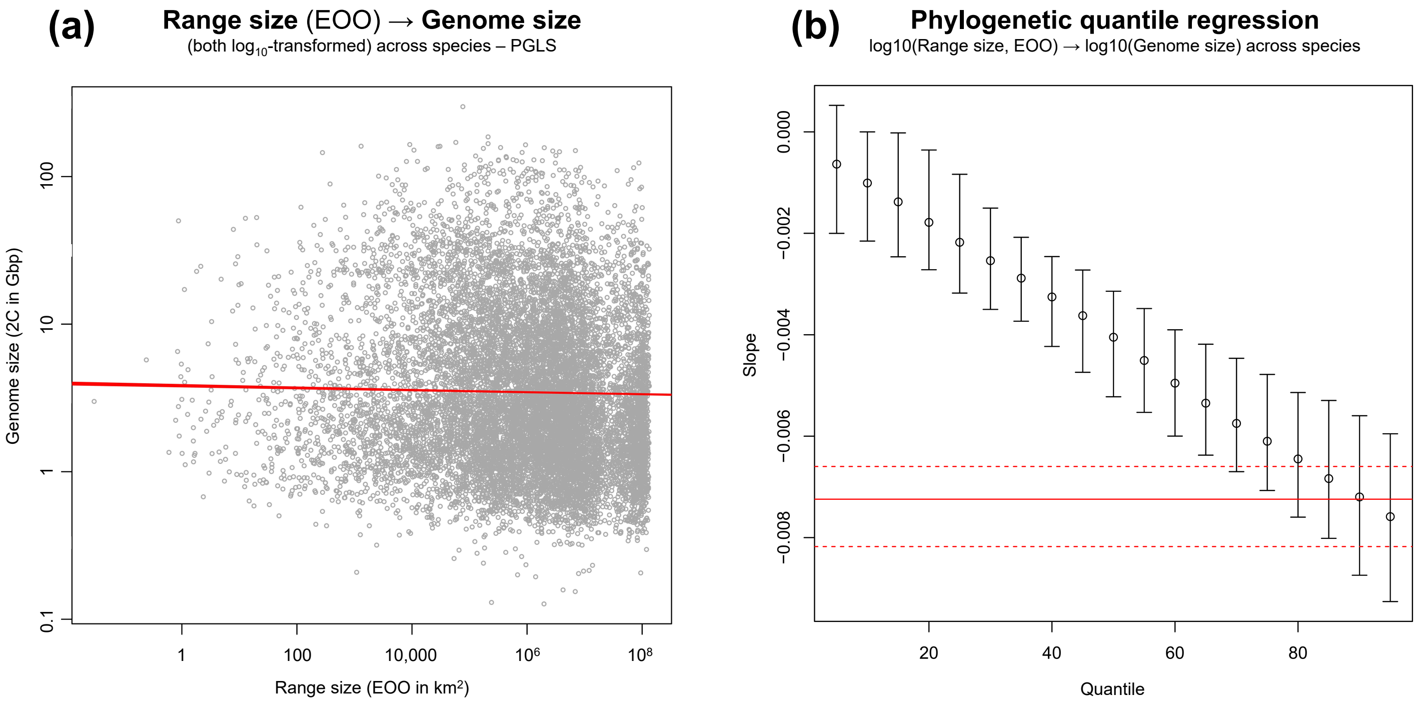


**For parallel analysis in which the number of occupied TDWGs was used for estimation of range size instead of the EOO (Extent of occurrence, based on GBIF data), see Fig. S5.**

**Fig. S3** Associations between genome and range size (as Extent of Occurence, EOO) per species considering phylogenetic relationships. The solid red line in (a) indicates the fit of the phylogenetic generalized least squares regression (PGLS), while the red line in (b) indicates the slope value from the phylogenetic quantile regression analysis. The slope estimates from the phylogenetic quantile regression, including 95% confidence intervals (error bars), are indicated in (b). Dashed red lines represent 95% confidence intervals. Both genome and range size are transformed by log10 in (a) and (b).

### Associations between genome and range size (no. of occupied TDWG regions)

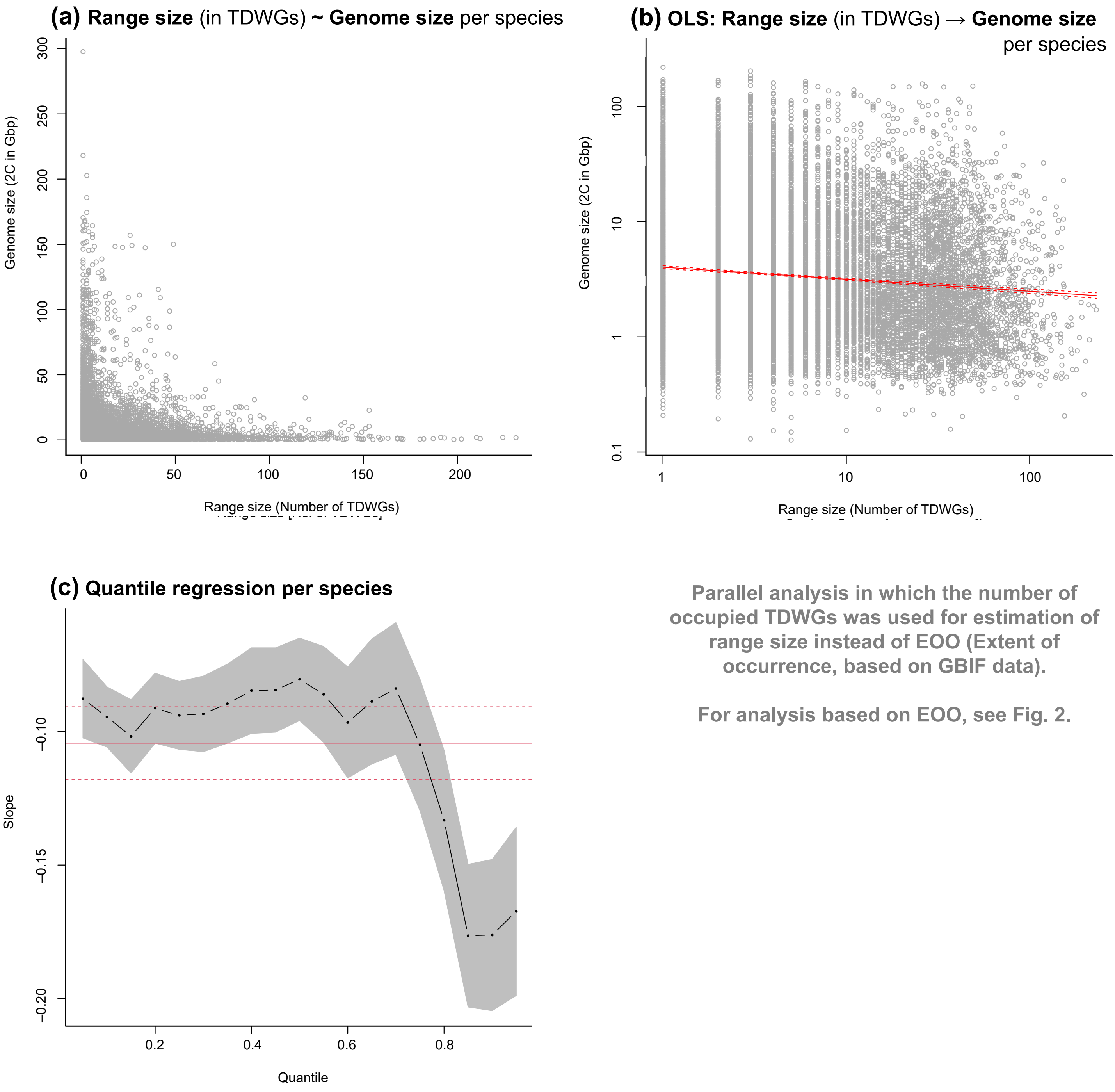


Parallel analysis in which the number of occupied TDWGs was used for estimation of range size instead of EOO (Extent of occurrence, based on GBIF data).

For analysis based on EOO, see Fig. 2.

**Fig. S4** Associations between genome and range size per species when the number of occupied TDWG regions (instead of Extent of Occurrence, EOO) is used as a measure of range size. The association of the raw data between genome and range size is shown in (a), whereas both variables are log-transformed in the other two plots (b,c). The slope estimates from the quantile regression, including 95% confidence intervals (dark grey), are indicated in (c). The solid red line in (b) indicates the fit of the ordinary least squares (OLS) regressions, while the solid red line in (c) indicates the slope value from the OLS analysis. Dashed red lines represent 95% confidence intervals.

### Associations between genome and range size (as no. of occupied TDWG regions) considering phylogenetic relationships

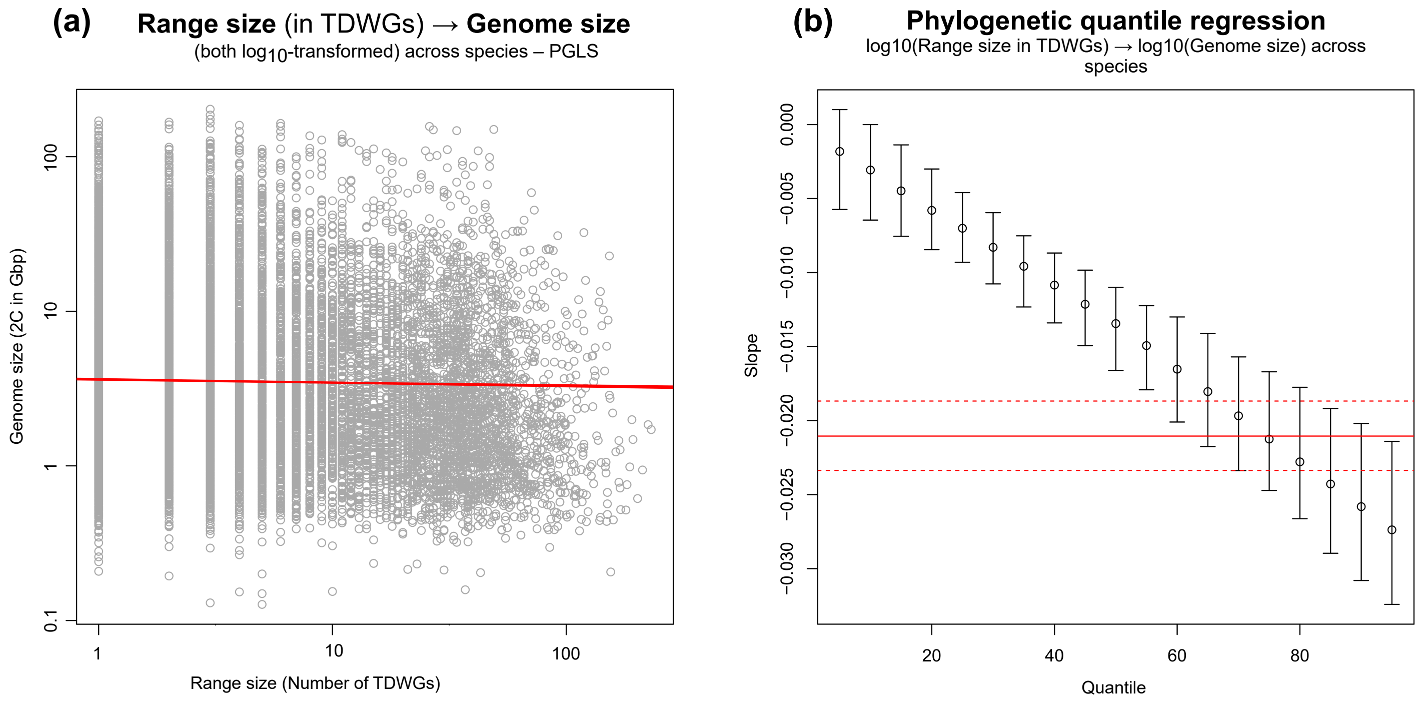


**For parallel analysis in which EOO (Extent of occurrence, based on GBIF data) was used for estimation of range size instead of the number of occupied TDWGs, see Fig. S3.**

**Fig. S5** Associations between genome and range size per species considering phylogenetic relationships when the number of occupied TDWG regions (instead of the Extent of Occurrence, EOO) is used as a measure of range size. The solid red line in (a) indicates the fit of the phylogenetic generalized least squares regression (PGLS), while the red line in (b) indicates the slope value from the phylogenetic quantile regression analysis. The slope estimates from the phylogenetic quantile regression, including 95% confidence intervals (error bars), are indicated in (b). Dashed red lines represent 95% confidence intervals. Both genome and range size are transformed by log10 in (a) and (b).

### Associations between mean chromosome size and range size (as Extent of Occurence, EOO)

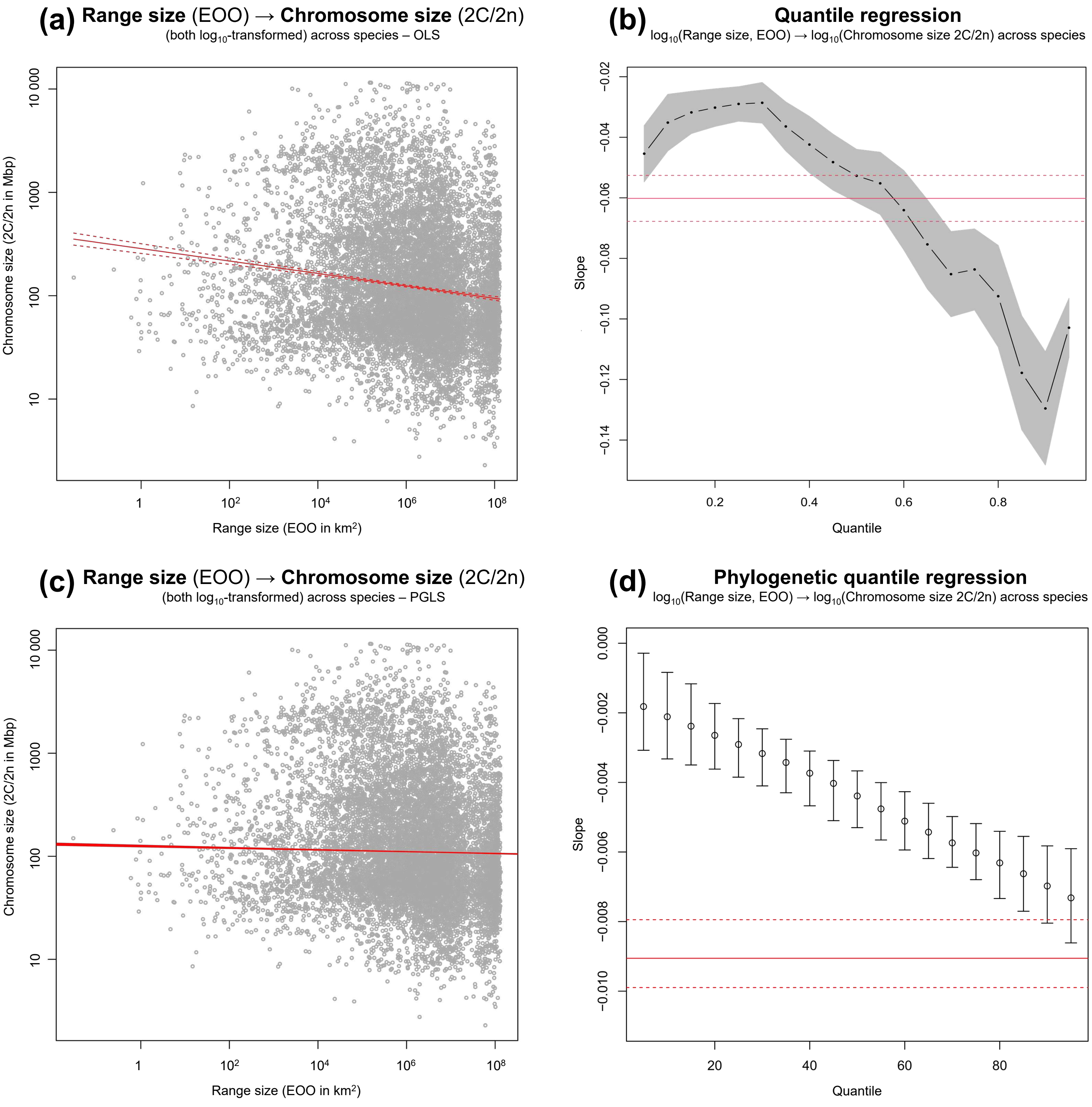


**Fig. S6** Associations between mean chromosome size and range size (as Extent of Occurence, EOO) per species. The solid red lines in (a) and (c) indicate the fit of the ordinary least squares (OLS) and phylogenetic generalized least squares (PGLS) regressions, respectively. The solid black circles and the gray shading in (b) represent the slope estimates and the 95% confidence intervals across 19 quantiles, whereas the hollow circles and the error bars in (d) indicate slope estimates and the 95% confidence intervals of the phylogenetic quantile regression. The horizontal red line in (b) represents the slope estimate of the OLS regression, while the horizontal red line in (d) shows the slope estimate of the PGLS regression. Dotted red lines in all four plots indicate the 95% confidence intervals of the slope estimates.

### Associations between mean chromosome size and range size (as no. of occupied TDWG regions)

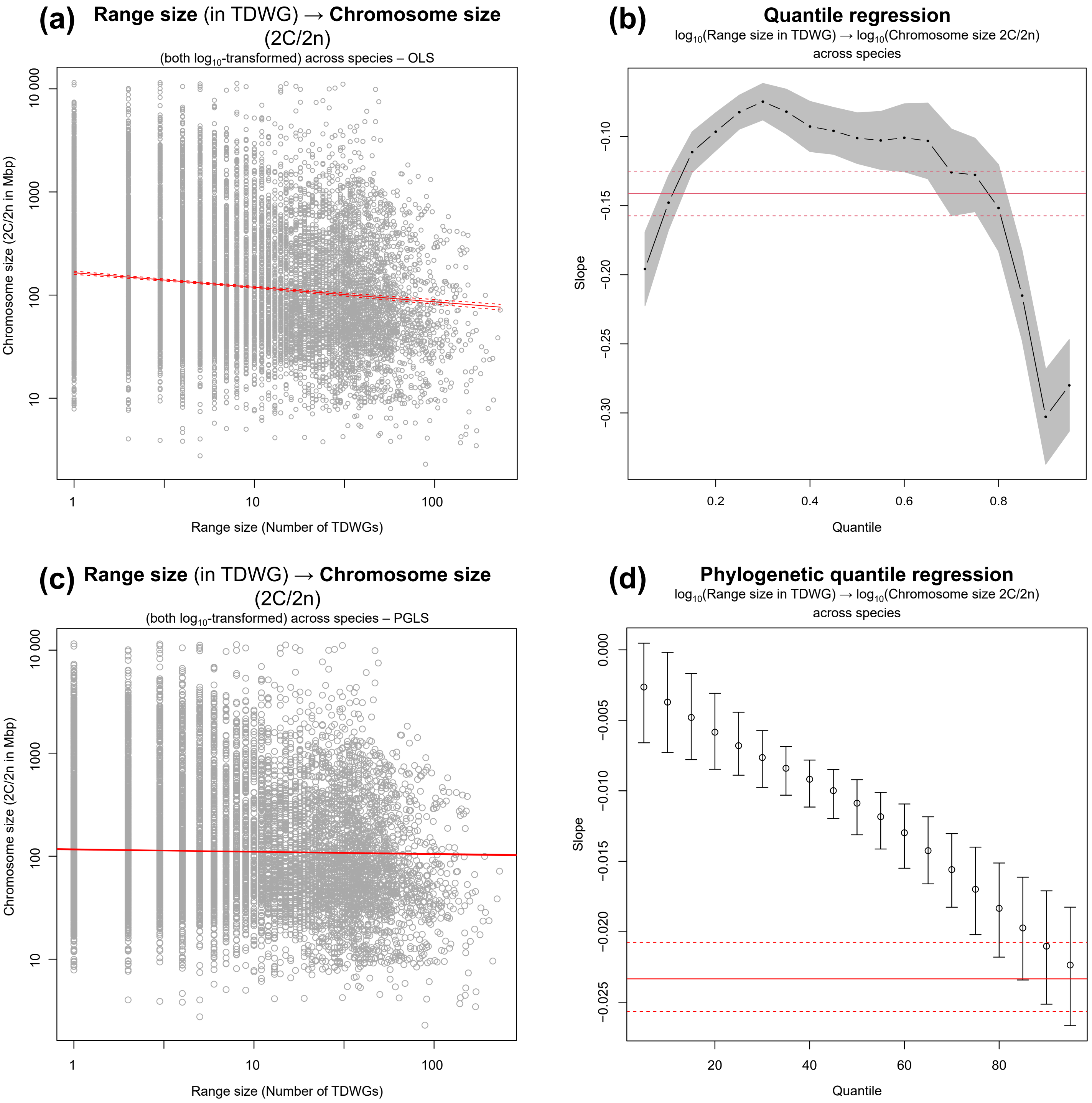


**Fig. S7** Associations between mean chromosome size and range size (as the number of occupied TDWG regions) per species. The solid red lines in (a) and (c) indicate the fit of the ordinary least squares (OLS) and phylogenetic generalized least squares (PGLS) regressions, respectively. The solid black circles and the gray shading in (b) represent the slope estimates and the 95% confidence intervals across 19 quantiles, whereas the hollow circles and the error bars in (d) indicate slope estimates and the 95% confidence intervals of the phylogenetic quantile regression. The horizontal red line in (b) represents the slope estimate of the OLS regression, while the horizontal red line in (d) shows the slope estimate of the PGLS regression. Dotted red lines in all four plots indicate the 95% confidence intervals of the slope estimates.

#### Distribution of mean genome size in the largest monocot and dicot families

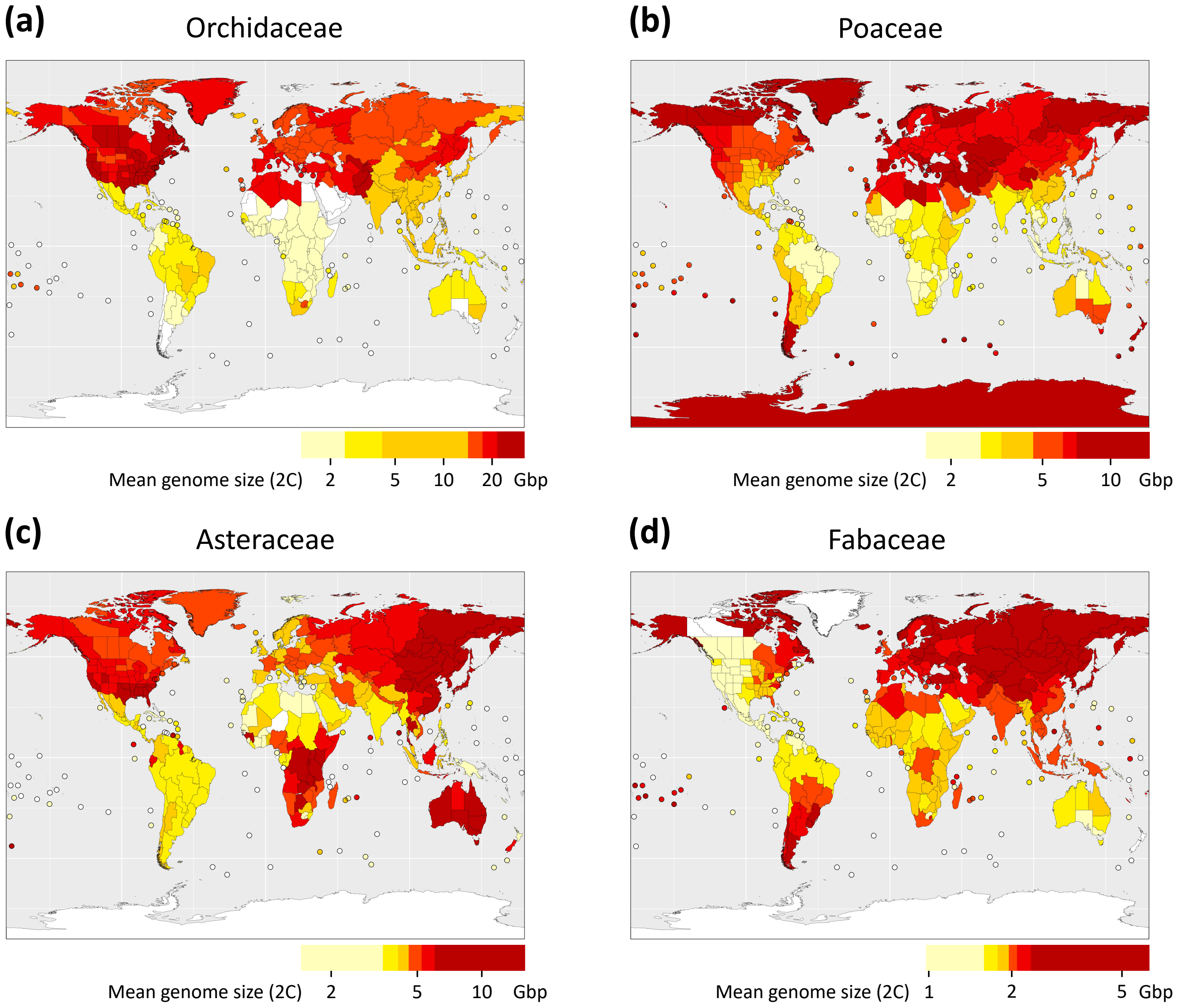


**Fig. S8** Mean genome sizes (2C; Gbp) averaged per TDWG region for the two most species-rich monocot (a – Orchidaceae, b – Poaceae) and dicot (c – Asteraceae, d – Fabaceae) families. Dark red colors indicate relatively large mean genome sizes, whereas light yellow shades indicate TDWG regions with relatively small mean genome sizes.

### Latitudinal gradient / distribution of mean chromosome size (2C/2n) in flowering plants

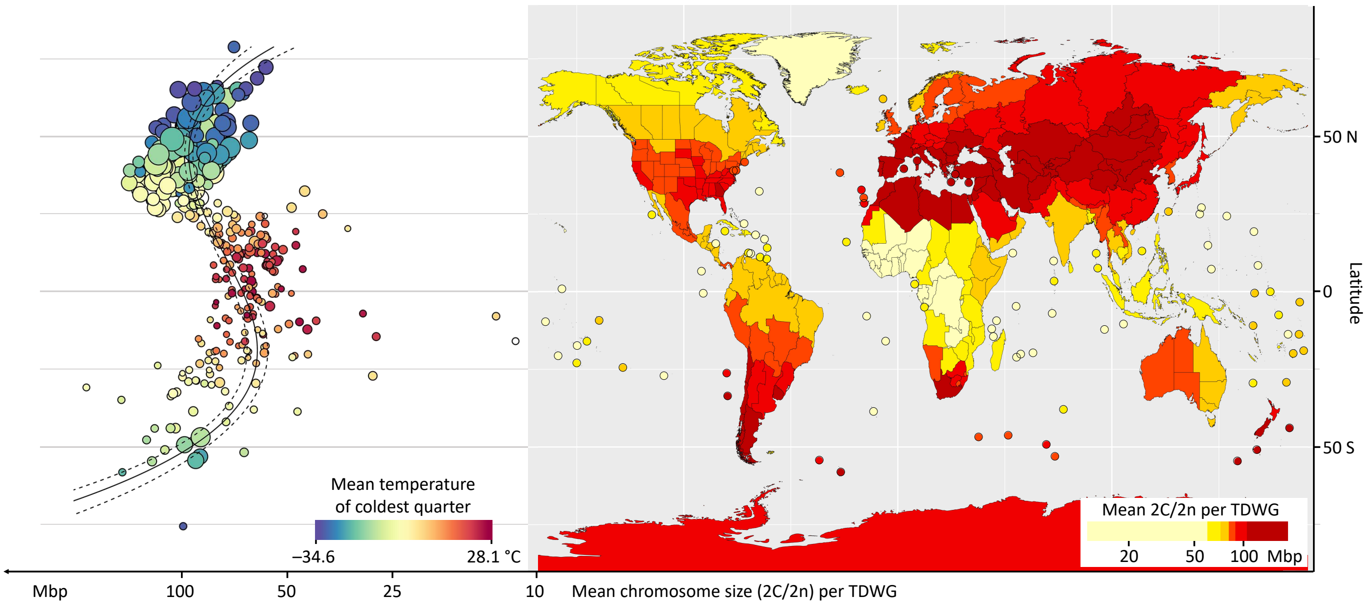


**Fig. S9** The global distribution of mean chromosome size in flowering plants calculated per TDWG region. The plot on the left side shows the distribution of mean chromosome sizes across latitudes, with dark reds indicating TDWG regions with high temperatures in the coldest quarter (BIO11 from Bioclim) and dark blues showing regions with low temperatures. The size of points in the plots indicates the weights used in the regression analysis (see Methods for details). The map on the right side shows the distribution of mean chromosome sizes mapped according to each TDWG region, where dark reds indicate relatively high values.

#### Latitudinal trends in mean genome sizes across Old World and New World

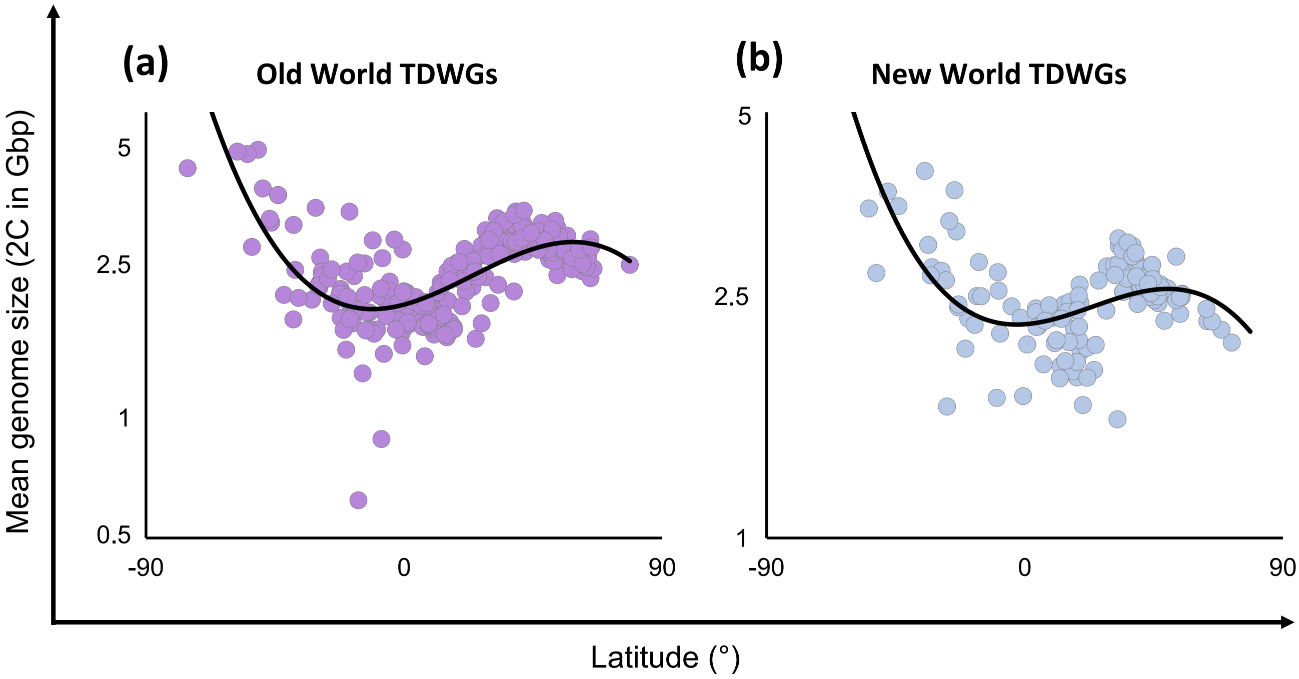


**Fig. S10** Mean genome sizes (2C; Gbp) across the global latitudinal gradient for the Old World (a) and New World (b). Circles in both plots represent the genome size averaged per TDWG region.

#### Latitudinal distribution of the percentage of growth forms in TDWG regions

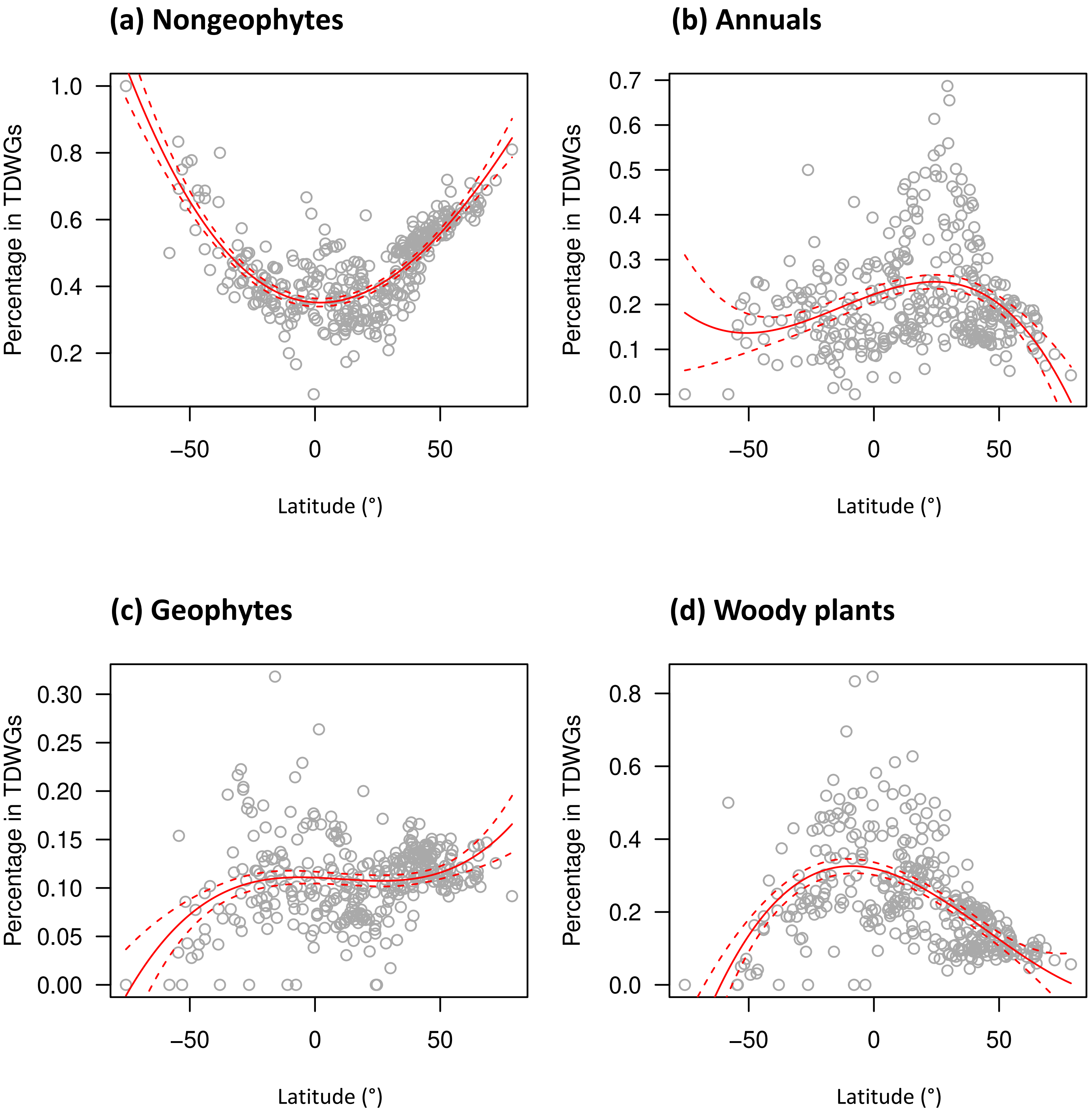


**Fig. S11** Latitudinal distribution of the percentage of (a) nongeophyte, (b) annual, (c) geophyte, and (d) woody species in our genome size dataset (Dataset S2).

#### Path analysis of causal relationships among temperature and growth forms (percentage) on the genome size distribution

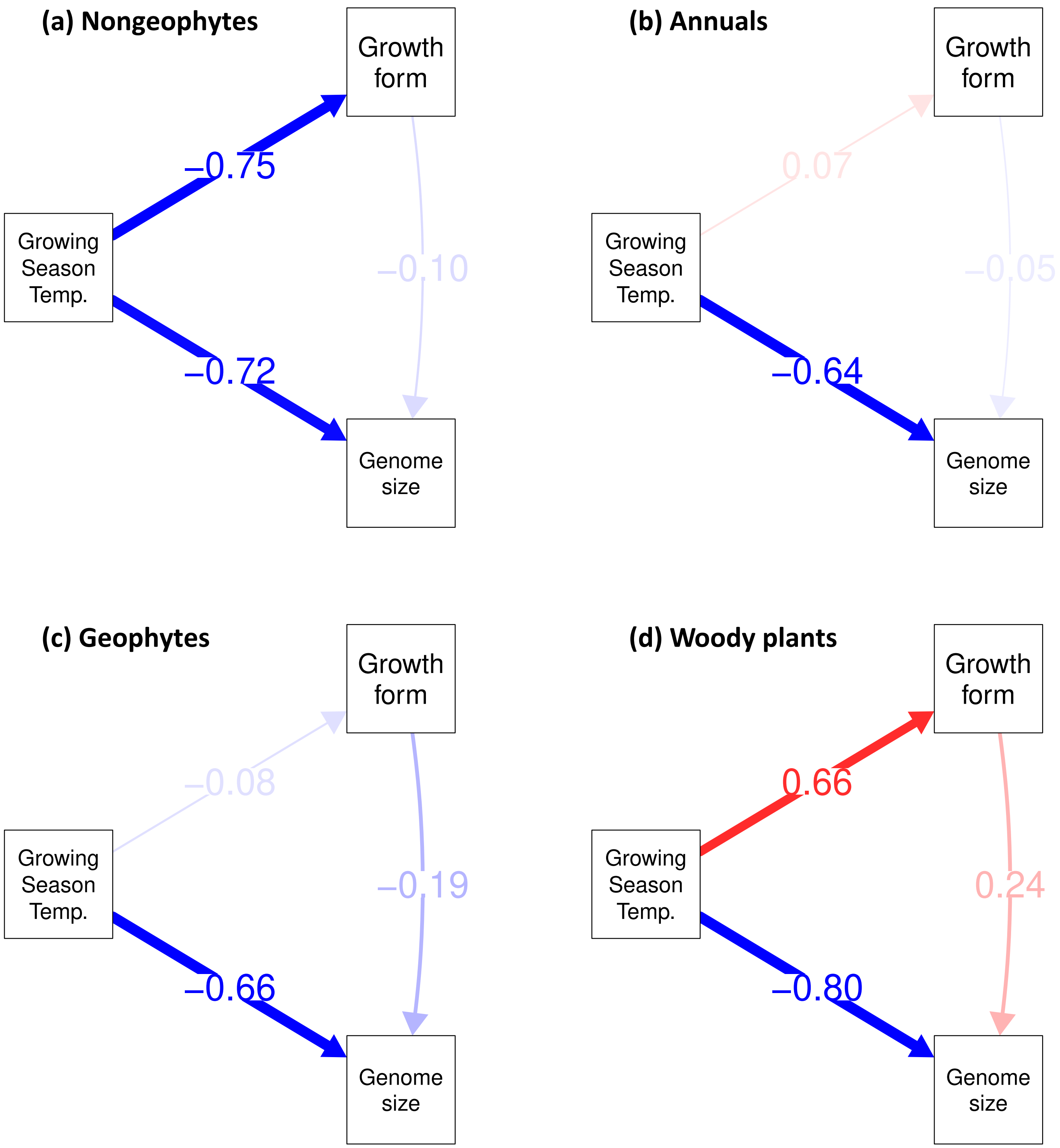


**Fig. S12** Path analysis of causal relationships among the effects of the growing season temperature (GST) and percentages of species of different growth forms on the mean genome size in TDWG regions: (a) nongeophytes, (b) annuals, (c) geophytes, and (d) woody species. The numbers indicate standardized regression coefficients from the path analyses. The arrows show the direction of the causal effects, their thickness indicates the relative effects, the fading indicates significance of the effect and the color indicates positive (red) or negative (blue) effect.

#### Latitudinal trends in mean genome sizes across glaciated and non-glaciated TDWGs

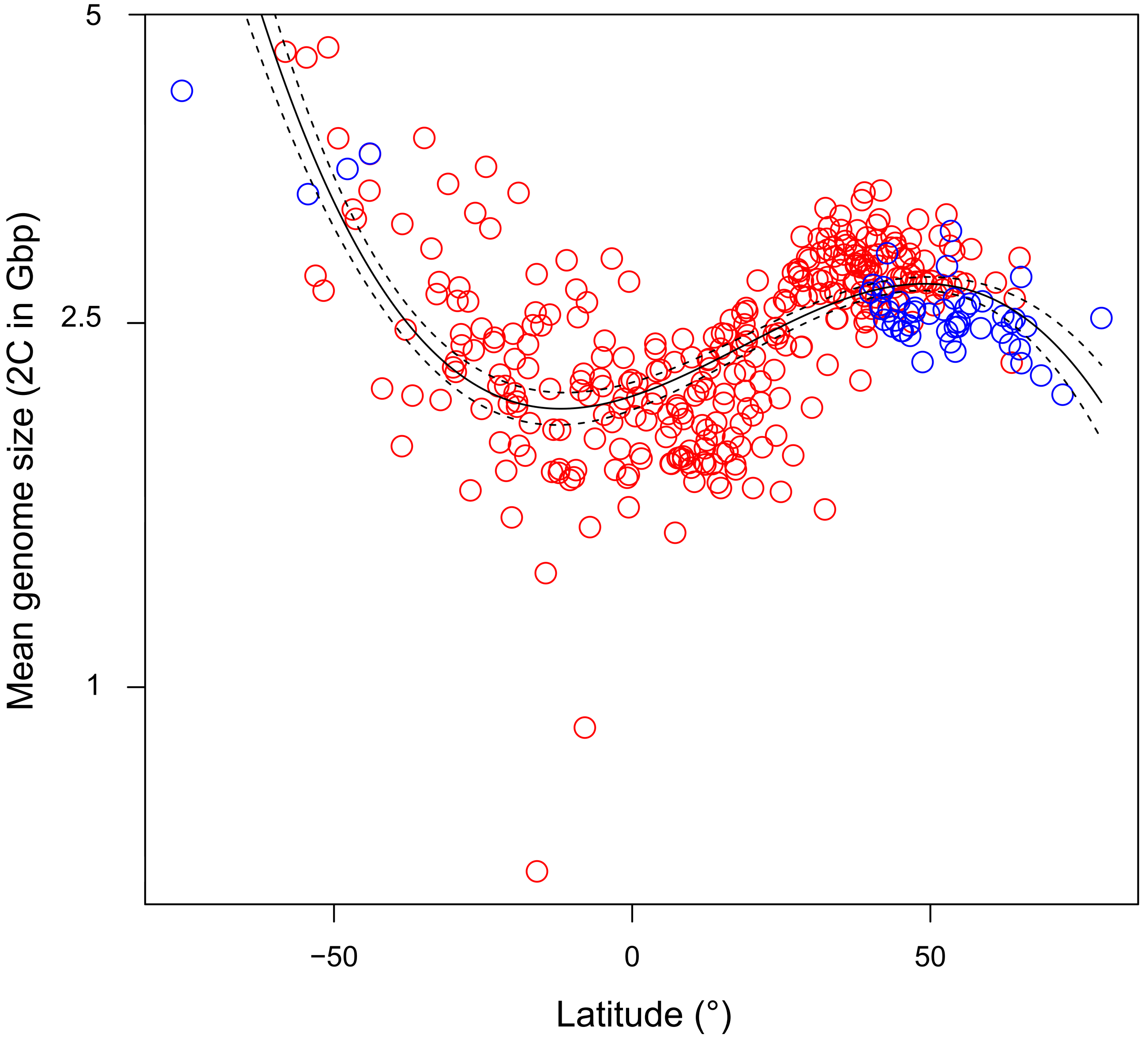


**Fig. S13** Mean genome sizes (2C; Gbp) across the global latitudinal gradient illustrating TDWG regions that were glaciated (blue) and non-glaciated (red) during the last glacial maximum (LGM) approximately 18,000 years before the present. We assessed the glaciation status of each TDWG region at the Last Glacial Maximum (LGM; ~18,000 years BP) using past climatic reconstructions from Ehlers (2015). We considered TDWG regions to be 'Glaciated' if their centroids were covered by the ice sheets during the LGM (Dataset S2).

*2. Supporting Information: Tables*

**Table S1: Bioclim variables as they explain the variance in 2C genome size across TDWGs in the polynomial regression of a given order**

| | Bioclim variable | N | order | $R^2_{adj}$ | explanation of the Bioclim variable |
|---|---|---|---|---|---|
| | BIO1 | 364 | 2 | 0.317 | Mean annual air temperature |
| | BIO3 | 364 | 3 | 0.314 | Isothermality |
| | BIO4 | 364 | 2 | 0.212 | Temperature seasonality |
| | BIO5 | 364 | 1 | 0.046 | Mean daily maximum air temperature of the warmest month |
| | BIO6 | 364 | 2 | 0.367 | Mean daily minimum air temperature of the coldest month |
| | BIO7 | 364 | 1 | 0.136 | Annual range of air temperature |
| | BIO8 | 364 | 2 | 0.326 | Mean daily mean air temperatures of the wettest quarter |
| | BIO9 | 364 | 2 | 0.086 | Mean daily mean air temperatures of the driest quarter |
| **Group 1** | BIO10 | 364 | 1 | 0.085 | Mean daily mean air temperatures of the warmest quarter |
| | BIO11 | 364 | 2 | 0.372 | Mean daily mean air temperatures of the coldest quarter |
| | GST | 365 | 2 | 0.408 | Mean temperature of the growing season |
| | NGD0 | 367 | 1 | 0.024 | Number of days at which mean daily air temperature > 0°C |
| | NGD5 | 363 | 2 | 0.157 | Number of days at which mean daily air temperature > 5°C |
| | NGS10 | 358 | 2 | 0.326 | Number of days at which mean daily air temperature > 10°C |
| | UVB1 | 367 | 2 | 0.346 | Annual Mean UV-B |
| | UVB3 | 367 | 3 | 0.145 | Mean UV-B of Highest Month |
| | UVB5 | 367 | 3 | 0.185 | Sum of Monthly Mean UV-B during Highest Quarter |
| **Group 2** | GDD0 | 367 | 1 | -0.002 | Growing degree days heat sum above 0°C |
| **Group 3** | BIO2 | 364 | 1 | 0.033 | Mean diurnal air temperature range |
| **Group 4** | BIO15 | 364 | 1 | 0.034 | Precipitation seasonality |
| **Group 5** | Aridity | 364 | 1 | 0.001 | Aridity index |
| | BIO12 | 364 | 1 | 0.066 | Annual precipitation amount |
| | BIO13 | 364 | 1 | 0.126 | Precipitation amount of the wettest month |
| | BIO14 | 364 | 1 | -0.003 | Precipitation amount of the driest month |
| **Group 6** | BIO16 | 364 | 1 | 0.118 | Mean monthly precipitation amount of the wettest quarter |
| | BIO17 | 364 | 1 | -0.002 | Mean monthly precipitation amount of the driest quarter |
| | BIO18 | 364 | 1 | 0.051 | Mean monthly precipitation amount of the warmest quarter |
| | BIO19 | 364 | 1 | 0.016 | Mean monthly precipitation amount of the coldest quarter |
| | GSL | 365 | 1 | 0.031 | Length of the growing season |

Variables were selected as predictors of 2C genome size in multiple regression based on the highest explained variance in their group. Aridity and GDD0 were not selected because of low explained variance. GST, BIO2, BIO13, and BIO15 were then included into a backward model selection process to select the best combination of variables predicting 2C genome size (see Methods for details).

**Table S2: Results of quantile regression of 2C genome size on range size (EOO)**

| Quantile regression model: log10(2C genome size) ~ log10(Range size) | | | | | |
|---|---|---|---|---|---|
| **Quantile** | **Model term** | **$b_i$** | **$se(b_i)$** | **t** | **P** |
| 5% | Intercept | 2.955 | 0.019 | 157.580 | 0 |
| | log10(Range size) | -0.024 | 0.003 | -7.533 | 0 |
| 10% | Intercept | 3.057 | 0.021 | 147.210 | 0 |
| | log10(Range size) | -0.025 | 0.003 | -7.503 | 0 |
| 15% | Intercept | 3.135 | 0.020 | 159.927 | 0 |
| | log10(Range size) | -0.024 | 0.003 | -7.339 | 0 |
| 20% | Intercept | 3.204 | 0.019 | 166.878 | 0 |
| | log10(Range size) | -0.024 | 0.003 | -7.546 | 0 |
| 25% | Intercept | 3.271 | 0.019 | 172.960 | 0 |
| | log10(Range size) | -0.025 | 0.003 | -8.126 | 0 |
| 30% | Intercept | 3.337 | 0.020 | 164.014 | 0 |
| | log10(Range size) | -0.027 | 0.003 | -8.153 | 0 |
| 35% | Intercept | 3.395 | 0.021 | 161.549 | 0 |
| | log10(Range size) | -0.027 | 0.003 | -7.817 | 0 |
| 40% | Intercept | 3.455 | 0.023 | 148.593 | 0 |
| | log10(Range size) | -0.027 | 0.004 | -7.013 | 0 |
| 45% | Intercept | 3.525 | 0.024 | 147.796 | 0 |
| | log10(Range size) | -0.028 | 0.004 | -7.113 | 0 |
| 50% | Intercept | 3.597 | 0.026 | 138.665 | 0 |
| | log10(Range size) | -0.030 | 0.004 | -7.092 | 0 |
| 55% | Intercept | 3.695 | 0.032 | 114.179 | 0 |
| | log10(Range size) | -0.035 | 0.005 | -6.958 | 0 |
| 60% | Intercept | 3.827 | 0.034 | 111.853 | 0 |
| | log10(Range size) | -0.044 | 0.005 | -8.151 | 0 |
| 65% | Intercept | 3.944 | 0.035 | 113.828 | 0 |
| | log10(Range size) | -0.049 | 0.006 | -8.883 | 0 |
| 70% | Intercept | 4.049 | 0.036 | 111.019 | 0 |
| | log10(Range size) | -0.051 | 0.006 | -8.765 | 0 |
| 75% | Intercept | 4.213 | 0.042 | 99.192 | 0 |
| | log10(Range size) | -0.061 | 0.007 | -9.283 | 0 |
| 80% | Intercept | 4.375 | 0.040 | 108.446 | 0 |
| | log10(Range size) | -0.069 | 0.006 | -10.762 | 0 |
| 85% | Intercept | 4.580 | 0.042 | 110.279 | 0 |
| | log10(Range size) | -0.081 | 0.007 | -11.957 | 0 |
| 90% | Intercept | 4.680 | 0.039 | 120.704 | 0 |
| | log10(Range size) | -0.071 | 0.007 | -10.870 | 0 |
| 95% | Intercept | 4.827 | 0.030 | 163.290 | 0 |
| | log10(Range size) | -0.060 | 0.005 | -11.347 | 0 |

$b_i$ – regression estimates of model terms.

**Table S3 Results of phylogenetic quantile regression of 2C genome size on range size (EOO)**

Phylogenetic quantile regression model: log10(2C genome size) ~ log10(Range size)

| Quantile | Model term | slope | L95%CI | U95%CI |
|---|---|---|---|---|
| 5% | log10(Range size) | -0.001 | -0.002 | 0.000 |
| 10% | log10(Range size) | -0.001 | -0.003 | 0.000 |
| 15% | log10(Range size) | -0.001 | -0.003 | 0.000 |
| 20% | log10(Range size) | -0.002 | -0.003 | 0.000 |
| 25% | log10(Range size) | -0.002 | -0.003 | -0.001 |
| 30% | log10(Range size) | -0.003 | -0.004 | -0.002 |
| 35% | log10(Range size) | -0.003 | -0.004 | -0.002 |
| 40% | log10(Range size) | -0.003 | -0.005 | -0.002 |
| 45% | log10(Range size) | -0.004 | -0.005 | -0.003 |
| 50% | log10(Range size) | -0.004 | -0.005 | -0.003 |
| 55% | log10(Range size) | -0.005 | -0.006 | -0.004 |
| 60% | log10(Range size) | -0.005 | -0.006 | -0.004 |
| 65% | log10(Range size) | -0.005 | -0.006 | -0.004 |
| 70% | log10(Range size) | -0.006 | -0.007 | -0.005 |
| 75% | log10(Range size) | -0.006 | -0.007 | -0.005 |
| 80% | log10(Range size) | -0.007 | -0.008 | -0.005 |
| 85% | log10(Range size) | -0.007 | -0.008 | -0.006 |
| 90% | log10(Range size) | -0.007 | -0.009 | -0.006 |
| 95% | log10(Range size) | -0.008 | -0.009 | -0.006 |

The intercept value is always zero because the phylogenetic quantile regression works with phylogenetic independent contrasts whose regression has to be forced through origin.

**Table S4: Results of quantile regression of genome size on range size (TDWGs)**

| Quantile regression model: log10(2C genome size) ~ log10(Range size) | | | | | |
|---|---|---|---|---|---|
| **Quantile** | **Model term** | **$b_i$** | **se($b_i$)** | **t** | **P** |
| 5% | Intercept | 2.899 | 0.007 | 412.607 | 0 |
| | log10(Range size) | -0.088 | 0.008 | -11.669 | 0 |
| 10% | Intercept | 3.011 | 0.006 | 492.020 | 0 |
| | log10(Range size) | -0.094 | 0.006 | -16.411 | 0 |
| 15% | Intercept | 3.087 | 0.006 | 495.326 | 0 |
| | log10(Range size) | -0.102 | 0.007 | -14.502 | 0 |
| 20% | Intercept | 3.146 | 0.005 | 584.281 | 0 |
| | log10(Range size) | -0.091 | 0.007 | -13.615 | 0 |
| 25% | Intercept | 3.208 | 0.007 | 482.003 | 0 |
| | log10(Range size) | -0.094 | 0.007 | -14.444 | 0 |
| 30% | Intercept | 3.266 | 0.006 | 510.241 | 0 |
| | log10(Range size) | -0.093 | 0.007 | -12.910 | 0 |
| 35% | Intercept | 3.319 | 0.007 | 509.107 | 0 |
| | log10(Range size) | -0.090 | 0.008 | -11.845 | 0 |
| 40% | Intercept | 3.374 | 0.007 | 460.997 | 0 |
| | log10(Range size) | -0.085 | 0.008 | -10.338 | 0 |
| 45% | Intercept | 3.438 | 0.007 | 472.582 | 0 |
| | log10(Range size) | -0.084 | 0.008 | -10.462 | 0 |
| 50% | Intercept | 3.491 | 0.007 | 475.151 | 0 |
| | log10(Range size) | -0.080 | 0.008 | -10.180 | 0 |
| 55% | Intercept | 3.557 | 0.009 | 385.882 | 0 |
| | log10(Range size) | -0.086 | 0.009 | -9.384 | 0 |
| 60% | Intercept | 3.644 | 0.010 | 360.321 | 0 |
| | log10(Range size) | -0.097 | 0.011 | -9.106 | 0 |
| 65% | Intercept | 3.720 | 0.010 | 355.561 | 0 |
| | log10(Range size) | -0.089 | 0.012 | -7.413 | 0 |
| 70% | Intercept | 3.815 | 0.012 | 318.872 | 0 |
| | log10(Range size) | -0.084 | 0.013 | -6.651 | 0 |
| 75% | Intercept | 3.934 | 0.013 | 298.276 | 0 |
| | log10(Range size) | -0.105 | 0.013 | -8.347 | 0 |
| 80% | Intercept | 4.077 | 0.015 | 272.455 | 0 |
| | log10(Range size) | -0.133 | 0.013 | -9.934 | 0 |
| 85% | Intercept | 4.253 | 0.015 | 290.314 | 0 |
| | log10(Range size) | -0.176 | 0.014 | -12.933 | 0 |
| 90% | Intercept | 4.411 | 0.013 | 336.729 | 0 |
| | log10(Range size) | -0.176 | 0.014 | -12.187 | 0 |
| 95% | Intercept | 4.607 | 0.016 | 296.578 | 0 |
| | log10(Range size) | -0.167 | 0.016 | -10.398 | 0 |

$b_i$ – regression estimates of model terms.

**Table S5: Results of phylogenetic quantile regression of genome size on range size (TDWGs)**

Phylogenetic quantile regression model: log10(2C genome size) ~ log10(Range size)

| Quantile | Model term | slope | L95%CI | U95%CI |
|---|---|---|---|---|
| 5% | log10(Range size) | -0.002 | -0.006 | 0.001 |
| 10% | log10(Range size) | -0.003 | -0.006 | 0.000 |
| 15% | log10(Range size) | -0.004 | -0.008 | -0.001 |
| 20% | log10(Range size) | -0.006 | -0.008 | -0.003 |
| 25% | log10(Range size) | -0.007 | -0.009 | -0.005 |
| 30% | log10(Range size) | -0.008 | -0.011 | -0.006 |
| 35% | log10(Range size) | -0.010 | -0.012 | -0.008 |
| 40% | log10(Range size) | -0.011 | -0.013 | -0.009 |
| 45% | log10(Range size) | -0.012 | -0.015 | -0.010 |
| 50% | log10(Range size) | -0.013 | -0.017 | -0.011 |
| 55% | log10(Range size) | -0.015 | -0.018 | -0.012 |
| 60% | log10(Range size) | -0.017 | -0.020 | -0.013 |
| 65% | log10(Range size) | -0.018 | -0.022 | -0.014 |
| 70% | log10(Range size) | -0.020 | -0.023 | -0.016 |
| 75% | log10(Range size) | -0.021 | -0.025 | -0.017 |
| 80% | log10(Range size) | -0.023 | -0.027 | -0.018 |
| 85% | log10(Range size) | -0.024 | -0.029 | -0.019 |
| 90% | log10(Range size) | -0.026 | -0.031 | -0.020 |
| 95% | log10(Range size) | -0.027 | -0.032 | -0.021 |

The intercept value is always zero because the phylogenetic quantile regression works with phylogenetic independent contrasts whose regression has to be forced through origin.

**Table S6: Results of OLS and PGLS regressions of mean chromosome size on range size (EOO)**

| | OLS model: log10(mean chromosome size) ~ log10(Range size) | | | | |
|---|---|---|---|---|---|
| **Model term** | **$b_i$** | **95%CI** | **t** | **P** | **$R^2_{adj}$** |
| **Intercept** | 2.456 | 2.409, 2.503 | 102.55 | <2E-16 | 0.022 |
| **log10(Range size)** | -0.060 | -0.068, -0.053 | -15.54 | <2E-16 | |

| | PGLS model: log10(mean chromosome size) ~ log10(Range size) | | | | |
|---|---|---|---|---|---|
| **Model term** | **$b_i$** | **95%CI** | **P** | **lambda** | **$R^2_{adj}$** |
| **Intercept** | 2.100 | 2.092, 2.105 | 4.02E-61 | 0.956 | 0.004 |
| **log10(Range size)** | -0.009 | -0.010, -0.008 | 2.08E-09 | | |

Results of ordinary least squares (OLS) and phylogenetic generalised least squares (PGLS) regression of mean

**Table S7: Results of quantile regression of mean chromosome size on range size (EOO)**

| Quantile regression model: log10(Mean chromosome size) ~ log10(Range size) | | | | | |
|---|---|---|---|---|---|
| **Quantile** | **Model term** | **$b_i$** | **se($b_i$)** | **t** | **P** |
| 5% | Intercept | 1.589 | 0.024 | 65.931 | 0 |
| | log10(Range size) | -0.045 | 0.005 | -9.576 | 0 |
| 10% | Intercept | 1.681 | 0.029 | 57.848 | 0 |
| | log10(Range size) | -0.035 | 0.005 | -7.365 | 0 |
| 15% | Intercept | 1.753 | 0.021 | 83.211 | 0 |
| | log10(Range size) | -0.032 | 0.004 | -8.924 | 0 |
| 20% | Intercept | 1.816 | 0.019 | 94.458 | 0 |
| | log10(Range size) | -0.030 | 0.003 | -9.535 | 0 |
| 25% | Intercept | 1.863 | 0.018 | 103.448 | 0 |
| | log10(Range size) | -0.029 | 0.003 | -9.957 | 0 |
| 30% | Intercept | 1.910 | 0.022 | 86.599 | 0 |
| | log10(Range size) | -0.029 | 0.003 | -8.346 | 0 |
| 35% | Intercept | 2.011 | 0.027 | 74.506 | 0 |
| | log10(Range size) | -0.036 | 0.004 | -8.802 | 0 |
| 40% | Intercept | 2.110 | 0.031 | 68.657 | 0 |
| | log10(Range size) | -0.042 | 0.005 | -8.912 | 0 |
| 45% | Intercept | 2.210 | 0.030 | 72.522 | 0 |
| | log10(Range size) | -0.048 | 0.005 | -10.205 | 0 |
| 50% | Intercept | 2.302 | 0.029 | 80.527 | 0 |
| | log10(Range size) | -0.053 | 0.004 | -11.811 | 0 |
| 55% | Intercept | 2.379 | 0.034 | 70.702 | 0 |
| | log10(Range size) | -0.055 | 0.005 | -10.520 | 0 |
| 60% | Intercept | 2.515 | 0.044 | 57.039 | 0 |
| | log10(Range size) | -0.064 | 0.007 | -9.587 | 0 |
| 65% | Intercept | 2.683 | 0.049 | 54.721 | 0 |
| | log10(Range size) | -0.075 | 0.008 | -10.026 | 0 |
| 70% | Intercept | 2.850 | 0.045 | 62.967 | 0 |
| | log10(Range size) | -0.085 | 0.007 | -11.857 | 0 |
| 75% | Intercept | 2.950 | 0.043 | 68.675 | 0 |
| | log10(Range size) | -0.084 | 0.007 | -12.230 | 0 |
| 80% | Intercept | 3.130 | 0.056 | 55.411 | 0 |
| | log10(Range size) | -0.093 | 0.009 | -10.788 | 0 |
| 85% | Intercept | 3.445 | 0.062 | 55.594 | 0 |
| | log10(Range size) | -0.118 | 0.010 | -12.277 | 0 |
| 90% | Intercept | 3.713 | 0.059 | 62.614 | 0 |
| | log10(Range size) | -0.130 | 0.010 | -13.579 | 0 |
| 95% | Intercept | 3.807 | 0.021 | 183.539 | 0 |
| | log10(Range size) | -0.103 | 0.005 | -20.790 | 0 |

$b_i$ – regression estimates of model terms.

**Table S8: Results of phylogenetic quantile regression of mean chromosome size on range size (EOO)**

| Phylogenetic quantile regression model: log10(mean chromosome size) ~ log10(Range size) | | | | |
|---|---|---|---|---|
| **Quantile** | **Model term** | **slope** | **L95%CI** | **U95%CI** |
| 5% | log10(Range size) | -0.002 | -0.003 | 0.000 |
| 10% | log10(Range size) | -0.002 | -0.003 | -0.001 |
| 15% | log10(Range size) | -0.002 | -0.003 | -0.001 |
| 20% | log10(Range size) | -0.003 | -0.004 | -0.002 |
| 25% | log10(Range size) | -0.003 | -0.004 | -0.002 |
| 30% | log10(Range size) | -0.003 | -0.004 | -0.002 |
| 35% | log10(Range size) | -0.003 | -0.004 | -0.003 |
| 40% | log10(Range size) | -0.004 | -0.005 | -0.003 |
| 45% | log10(Range size) | -0.004 | -0.005 | -0.003 |
| 50% | log10(Range size) | -0.004 | -0.005 | -0.004 |
| 55% | log10(Range size) | -0.005 | -0.006 | -0.004 |
| 60% | log10(Range size) | -0.005 | -0.006 | -0.004 |
| 65% | log10(Range size) | -0.005 | -0.006 | -0.004 |
| 70% | log10(Range size) | -0.006 | -0.006 | -0.005 |
| 75% | log10(Range size) | -0.006 | -0.007 | -0.005 |
| 80% | log10(Range size) | -0.006 | -0.007 | -0.005 |
| 85% | log10(Range size) | -0.007 | -0.008 | -0.005 |
| 90% | log10(Range size) | -0.007 | -0.008 | -0.005 |
| 95% | log10(Range size) | -0.007 | -0.009 | -0.006 |

The intercept value is always zero because the phylogenetic quantile regression works with phylogenetic independent contrasts whose regression has to be forced through origin.

**Table S9: Results of OLS and PGLS regressions of mean chromosome size on range size (TDWGs)**

| | OLS model: log10(mean chromosome size) ~ log10(Range size) | | | | |
|---|---|---|---|---|---|
| **Model term** | **$b_i$** | **95%CI** | **t** | **P** | **$R^2_{adj}$** |
| **Intercept** | 2.217 | 2.201, 2.232 | 278.40 | <2e-16 | 0.021 |
| **log10(Range size)** | -0.141 | -0.157, -0.125 | -17.14 | <2e-16 | |

| | PGLS model: log10(mean chromosome size) ~ log10(Range size) | | | | |
|---|---|---|---|---|---|
| **Model term** | **$b_i$** | **95%CI** | **P** | **lambda** | **$R^2_{adj}$** |
| **Intercept** | 2.066 | 2.066, 2.067 | 1.57E-58 | 0.962 | 0.004 |
| **log10(Range size)** | -0.023 | -0.024, -0.023 | 4.32E-09 | | |

Results of ordinary least squares (OLS) and phylogenetic generalised least squares (PGLS) regression of mean chromosome size on range size. $b_i$ - regression estimates of model terms; 95%CI - lower and upper 95% confidence intervals of the regression estimates; $R^2_{adj}$ - R squared adjusted indicating explained variance. The OLS analysis was perfromed with 13,996 species. The PGLS analysis was perfromed with 10,670 species.  The PGLS was performed repeatedly with one hundred different trees (see Methods). Therefore, the values for PGLS are averages across these one hundred regressions.

**Table S10: Results of quantile regression of mean chromosome size on range size (TDWGs)**

| Quantile regression model: log10(mean chromosome size) ~ log10(Range size) | | | | | |
|---|---|---|---|---|---|
| **Quantile** | **Model term** | **$b_i$** | **$se(b_i)$** | **t** | **P** |
| 5% | Intercept | 1.491 | 0.010 | 152.814 | 0 |
| | log10(Range size) | -0.196 | 0.014 | -14.364 | 0 |
| 10% | Intercept | 1.601 | 0.007 | 240.353 | 0 |
| | log10(Range size) | -0.148 | 0.010 | -14.692 | 0 |
| 15% | Intercept | 1.659 | 0.006 | 268.413 | 0 |
| | log10(Range size) | -0.111 | 0.008 | -14.814 | 0 |
| 20% | Intercept | 1.713 | 0.006 | 299.498 | 0 |
| | log10(Range size) | -0.097 | 0.007 | -13.752 | 0 |
| 25% | Intercept | 1.754 | 0.005 | 344.908 | 0 |
| | log10(Range size) | -0.082 | 0.006 | -13.073 | 0 |
| 30% | Intercept | 1.797 | 0.007 | 268.692 | 0 |
| | log10(Range size) | -0.075 | 0.007 | -11.056 | 0 |
| 35% | Intercept | 1.857 | 0.009 | 213.299 | 0 |
| | log10(Range size) | -0.082 | 0.008 | -9.853 | 0 |
| 40% | Intercept | 1.932 | 0.010 | 202.680 | 0 |
| | log10(Range size) | -0.093 | 0.009 | -9.981 | 0 |
| 45% | Intercept | 2.000 | 0.009 | 229.814 | 0 |
| | log10(Range size) | -0.096 | 0.009 | -11.055 | 0 |
| 50% | Intercept | 2.065 | 0.010 | 216.425 | 0 |
| | log10(Range size) | -0.101 | 0.009 | -10.771 | 0 |
| 55% | Intercept | 2.136 | 0.010 | 206.850 | 0 |
| | log10(Range size) | -0.103 | 0.011 | -9.590 | 0 |
| 60% | Intercept | 2.216 | 0.012 | 179.613 | 0 |
| | log10(Range size) | -0.101 | 0.013 | -8.028 | 0 |
| 65% | Intercept | 2.314 | 0.015 | 158.329 | 0 |
| | log10(Range size) | -0.103 | 0.014 | -7.370 | 0 |
| 70% | Intercept | 2.450 | 0.016 | 150.790 | 0 |
| | log10(Range size) | -0.126 | 0.016 | -7.864 | 0 |
| 75% | Intercept | 2.567 | 0.013 | 193.387 | 0 |
| | log10(Range size) | -0.128 | 0.014 | -9.406 | 0 |
| 80% | Intercept | 2.714 | 0.019 | 139.610 | 0 |
| | log10(Range size) | -0.152 | 0.016 | -9.471 | 0 |
| 85% | Intercept | 2.925 | 0.021 | 142.058 | 0 |
| | log10(Range size) | -0.215 | 0.017 | -12.974 | 0 |
| 90% | Intercept | 3.214 | 0.019 | 168.460 | 0 |
| | log10(Range size) | -0.303 | 0.018 | -17.297 | 0 |
| 95% | Intercept | 3.424 | 0.013 | 257.932 | 0 |
| | log10(Range size) | -0.280 | 0.017 | -16.527 | 0 |

**Table S11: Results of phylogenetic quantile regression of mean chromosome size on range size (TDWGs)**

Phylogenetic quantile regression model: log10(2C genome size) ~ log10(Range size)

| Quantile | Model term | slope | L95%CI | U95%CI |
|---|---|---|---|---|
| 5% | log10(Range size) | -0.003 | -0.007 | 0.000 |
| 10% | log10(Range size) | -0.004 | -0.007 | 0.000 |
| 15% | log10(Range size) | -0.005 | -0.008 | -0.002 |
| 20% | log10(Range size) | -0.006 | -0.008 | -0.003 |
| 25% | log10(Range size) | -0.007 | -0.009 | -0.004 |
| 30% | log10(Range size) | -0.008 | -0.010 | -0.006 |
| 35% | log10(Range size) | -0.008 | -0.010 | -0.007 |
| 40% | log10(Range size) | -0.009 | -0.011 | -0.008 |
| 45% | log10(Range size) | -0.010 | -0.012 | -0.008 |
| 50% | log10(Range size) | -0.011 | -0.013 | -0.009 |
| 55% | log10(Range size) | -0.012 | -0.014 | -0.010 |
| 60% | log10(Range size) | -0.013 | -0.015 | -0.011 |
| 65% | log10(Range size) | -0.014 | -0.017 | -0.012 |
| 70% | log10(Range size) | -0.016 | -0.018 | -0.013 |
| 75% | log10(Range size) | -0.017 | -0.020 | -0.014 |
| 80% | log10(Range size) | -0.018 | -0.022 | -0.015 |
| 85% | log10(Range size) | -0.020 | -0.023 | -0.016 |
| 90% | log10(Range size) | -0.021 | -0.025 | -0.017 |
| 95% | log10(Range size) | -0.022 | -0.027 | -0.018 |

The intercept value is always zero because the phylogenetic quantile regression works with phylogenetic independent contrasts whose regression has to be forced through origin.

**Table S12: Additional regressions of 2C genome size on other biologically relevant variables**

| Model term | $b_i$ | 95%CI | t | P | $R^2_{adj}$ |
|---|---|---|---|---|---|
| Polynomial regression (N=367): log10(2C genome size) ~ UVB1 + UVB1$^2$ | | | | | |
| Intercept | 3.414 | <3.361, 3.467> | 127.37 | <2E-16 | |
| UVB1 | 3.89E-05 | <5.95E-06, 7.18E-05> | 2.322 | 0.0208 | 0.3457 |
| UVB1$^2$ | -1.03E-08 | <-1.48E-08, -5.84E-09> | -4.533 | <8E-06 | |
| Polynomial regression (N=362): log10(2C genome size) ~ GST + GST$^2$ + BIO11 + BIO11$^2$ | | | | | |
| Intercept | 3.508 | <3.455, 3.562> | 128.493 | <2E-16 | |
| GST | -2.52E-03 | <-9.0499e-03, 4.0133e-03> | -0.758 | 0.4488 | |
| GST$^2$ | -1.76E-04 | <-3.8406e-04, 3.1972e-05> | -1.664 | 0.0969 | 0.4635 |
| BIO11 | 3.94E-04 | <-6.5567e-04, 1.4439e-03> | 0.738 | 0.4608 | |
| BIO11$^2$ | -4.99E-05 | <-9.9424e-05, -4.4982e-07> | -1.985 | 0.048 | |
| Polynomial regression (N=365): log10(2C genome size) ~ GST + GST$^2$ + log10(range size) | | | | | |
| Intercept | 3.854 | <3.7182, 3.9906> | 55.661 | <2E-16 | |
| GST | -1.13E-03 | <-0.0077, 0.0054> | -0.341 | 0.733 | 0.4614 |
| GST$^2$ | -2.00E-04 | <-0.0004, 0> | -2.146 | 3.25E-02 | |
| log10(range size) | -0.05253 | <-0.0708, -0.0342> | -5.64 | 3.45E-08 | |
| Polynomial regression (N=365): log10(2C genome size) ~ GSL | | | | | |
| Intercept | 3.426 | <3.401, 3.450> | 272.35 | <2E-16 | 0.0311 |
| GSL | -1.62E-04 | <-2.52E-04, -7.62E-05> | -3.561 | 0.0004 | |
| Polynomial regression (N=47): log10(2C genome size) ~ GSL [only for TDGWs from 48.93° northward] | | | | | |
| Intercept | 3.413 | <3.386, 3.440> | 256.048 | <2E-16 | -0.0109 |
| GSL | 4.94E-05 | <-9.06E-05, 1.89E-04> | 0.711 | 0.4810 | |

UVB1 - mean annual UVB; GST - mean temperature of growing season; range size - range size of species presented in TDWG for which we had 2C genome size; GSL - mean length of the growing season; BIO11 - Daily mean air temperatures of the coldest quarter; bi - regression estimates of model terms; 95%CI - lower and upper 95% confidence intervals of the regression estimates; $R^2_{adj}$ - R squared adjusted indicating explained variance.

**Table S13: Results of regressions of 2C genome size on percentage of growth forms in TDWGs**

model: log10(2C genome size) ~ percentage of nongeophytes

| Model term | $b_i$ | 95%CI | t | P | $R^2_{adj}$ |
|---|---|---|---|---|---|
| **Intercept** | 3.255 | <3.2211, 3.2883> | 190.50 | <2E-16 | 0.211 |
| **% of nongeophytes** | 0.321 | <0.2573, 0.3842> | 9.94 | <2E-16 | |

model: log10(2C genome size) ~ percentage of annuals

| Model term | $b_i$ | 95%CI | t | P | $R^2_{adj}$ |
|---|---|---|---|---|---|
| **Intercept** | 3.444 | <3.4236, 3.4640> | 334.65 | <2E-16 | 0.019 |
| **% of annuals** | -0.119 | <-0.2014, -0.0356> | -2.81 | 0.00519 | |

model: log10(2C genome size) ~ percentage of geophytes

| Model term | $b_i$ | 95%CI | t | P | $R^2_{adj}$ |
|---|---|---|---|---|---|
| **Intercept** | 3.494 | <3.4675, 3.5212> | 256.03 | <2E-16 | 0.086 |
| **% of geophytes** | -0.702 | <-0.9336, -0.4694> | -5.94 | 6.54E-09 | |

model: log10(2C genome size) ~ percentage of woody plants

| Model term | $b_i$ | 95%CI | t | P | $R^2_{adj}$ |
|---|---|---|---|---|---|
| **Intercept** | 3.456 | <3.4396, 3.4717> | 423.19 | <2E-16 | 0.078 |
| **% of woody plants** | -0.221 | <-0.2974, -0.1440> | -5.66 | 3.10E-08 | |

$b_i$ - regression estimates of model terms; 95%CI - lower and upper 95% confidence intervals of the regression estimates; $R^2_{adj}$ - R squared adjusted indicating explained variance. The analyses were perfromed with 367 TDWGs.

**Table S14: Results of regressions of 2C genome size on additive effects of GST and percentage of growth forms in TDWGs**

model: log10(2C genome size) ~ GST + $GST^2$ + percentage of nongeophytes

| Model term | $b_i$ | 95%CI | t | P | $R^2_{adj}$ | increase in $R^2_{adj}$ |
|---|---|---|---|---|---|---|
| **Intercept** | 3.5630 | <3.4810, 3.6455> | 85.19 | <2e-16 | 0.408 | NA |
| **GST** | -0.0032 | <-0.0085, 0.0022> | -1.17 | 0.2436 | | |
| **$GST^2$** | -0.0002 | <-0.0004, -0.0001> | -2.97 | 0.0032 | | |
| **% of nongeophytes** | -0.0883 | <-0.1800, 0.0034> | -1.89 | 0.0592 | | |

model: log10(2C genome size) ~ GST + $GST^2$ + percentage of annuals

| Model term | $b_i$ | 95%CI | t | P | $R^2_{adj}$ | increase in $R^2_{adj}$ |
|---|---|---|---|---|---|---|
| **Intercept** | 3.4910 | <3.4531, 3.5295> | 179.74 | <2E-16 | 0.402 | NA |
| **GST** | -0.0012 | <-0.0062, 0.0037> | -0.48 | 0.6292 | | |
| **$GST^2$** | -0.0002 | <-0.0004, -0.0001> | -3.18 | 0.0016 | | |
| **% of annuals** | 0.0118 | <-0.0519, 0.0756> | 0.37 | 0.7157 | | |

model: log10(2C genome size) ~ GST + $GST^2$ + percentage of geophytes

| Model term | $b_i$ | 95%CI | t | P | $R^2_{adj}$ | increase in $R^2_{adj}$ |
|---|---|---|---|---|---|---|
| **Intercept** | 3.5190 | <3.4811, 3.5560> | 184.55 | <2E-16 | 0.442 | 0.034 |
| **GST** | 0.0029 | <-0.0021, 0.0079> | 1.15 | 0.2520 | | |
| **$GST^2$** | -0.0004 | <-0.0005, -0.0002> | -4.74 | 3.05E-06 | | |
| **% of geophytes** | -0.4947 | <-0.6857, -0.3037> | -5.09 | 5.66E-07 | | |

model: log10(2C genome size) ~ percentage of woody plants

| Model term | $b_i$ | 95%CI | t | P | $R^2_{adj}$ | increase in $R^2_{adj}$ |
|---|---|---|---|---|---|---|
| **Intercept** | 3.4890 | <3.4518, 3.5256> | 186.05 | <2E-16 | 0.421 | 0.014 |
| **GST** | -0.0018 | <-0.0066, 0.0030> | -0.73 | 0.4671 | | |
| **$GST^2$** | -0.0003 | <-0.0004, -0.0001> | -3.80 | 0.0002 | | |
| **% of woody plants** | 0.1490 | <0.0646, 0.2335> | 3.47 | 0.0006 | | |

$b_i$ - regression estimates of model terms; 95%CI - lower and upper 95% confidence intervals of the regression estimates; t - t statistics; P - significance; $R^2_{adj}$ - R squared adjusted indicating explained variance. The analyses were perfromed with 365 TDWGs. The increase in $R^2_{adj}$ after adding the percentage of growth forms into a model was not applicable for nongeophytes and annuals as their effect on genome size was insignificant (see their P-values). The GST alone explained 40.75 % (Table 2).
